# Relationships between phenotypic differences in punishment behaviour and prefrontal cortex network engagement

**DOI:** 10.1101/2025.10.24.684296

**Authors:** Min Lou, Saraa Al-Saddik, Michelle H. Shen, Luke J. Keevers, Philip Jean-Richard-dit-Bressel

**Author notes:** **Corresponding author:** Philip Jean-Richard-dit-Bressel.

## Abstract

Avoiding actions with negative consequences is fundamental to adaptive behaviour, yet individuals differ drastically in their tendencies to avoid to punishment. Using a conditioned punishment task in male and female rats, we replicate findings of a bimodal distribution of punishment avoidance, and identify four distinct behavioural phenotypes (Sensitive, Hypersensitive, Insensitive, Generalised) that differed in their punishment avoidance, unpunished reward-seeking, and Pavlovian fear. We examined how these phenotypic differences related to c-Fos (transcriptional activity marker) across prefrontal cortex (PFC) regions. While region-level c-Fos differences were minimal, covariance and graph-theoretic analyses revealed three functional PFC networks predominantly composed of medial PFC, ventrolateral PFC, or sensorimotor regions. Activity in the ventrolateral PFC network predicted punishment avoidance, while activity in the medial PFC network predicted generalised suppression during initial punishment learning. These findings suggest phenotypic differences in punishment sensitivity is underpinned by differential recruitment of distinct cortical networks that drive adaptive versus maladaptive responses to negative outcomes.

---

Animals are generally sensitive to the consequences of their actions, and will dynamically adjust their behaviour to better obtain rewards and avoid harm. This adaptive tendency to increase rewarded behaviours and reduce detrimental ones (known as reinforcement and punishment, respectively) is crucial to survival and thriving, and is unsurprisingly preserved across species (Thorndike, 1911). However, it has been shown that individuals differ drastically in their ability to suppress harmful behaviours; some individuals are punishment-sensitive and selectively suppress punished actions while others are punishment-insensitive and persist despite ongoing detriment (B. T. Chen et al., 2013; Y. Chen et al., 2022; Domi et al., 2021; Engeln & Ahmed, 2024; Gaetani et al., 2025; Jean-Richard-dit-Bressel et al., 2019, 2021, 2023; Marchant et al., 2018; McDonald et al., 2024; Zeng et al., 2025). Beyond its relevance to everyday decision-making, differences in punishment sensitivity are implicated in various clinical conditions. Insensitivity to punishment is implicated in substance use disorders (McNally et al., 2023; Pelloux et al., 2007; Torres et al., 2017; Vanderschuren et al., 2017), gambling addiction (De Ruiter et al., 2009), conduct disorder (Dadds & Salmon, 2003; Elster et al., 2024), and obsessive-compulsive disorder (Figee et al., 2016; Palminteri et al., 2012). Conversely, hypersensitivity to negative outcomes is a hallmark of anxiety (Broman-Fulks et al., 2014; Giorgetta et al., 2012), depression (Eshel & Roiser, 2010; Hevey et al., 2017), and eating disorders (Harrison et al., 2010).

A barrier to understanding mechanisms of punishment insensitivity is that the specific psychological constructs driving differences in punishment behaviour can be ambiguous. Punishment insensitivity can stem from differences in how punishing and/or rewarding outcomes are valued (low aversive motivation [Corr, 2004; Engeln & Ahmed, 2024] vs. reward dominance accounts [O’Brien & Frick, 1996; Robinson & Berridge, 2003]). Alternatively, insensitivity can result from impaired contingency learning, where individuals fail to accurately identify the cause of punishment, and instead fall into learning traps (Maier & Jackson, 1979; Maier & Seligman, 2016; Rich & Gureckis, 2018; Seligman, 1972).

Addressing this, we recently leveraged a conditioned punishment task (Killcross et al., 1997) to disentangle motivational and associative underpinnings of inter-individual differences in avoidance (Jean-Richard-dit-Bressel et al., 2019). In this task, animals can press two levers (R1, R2) for food reward, but R1 responses come to occasion an aversive cue (CS+) that precipitates footshock, while R2 presses occasion a neutral cue (CS-) without further consequence. When subjecting 48 animals to this task, avoidance of punished R1 relative to unpunished R2 was bimodal across subjects. Data-driven clustering identified 3 distinct profiles of behaviour in the task: 1) a “punishment-sensitive” profile that showed strong selective avoidance of R1, 2) an “punishment-insensitive” profile of modest, equivalent suppression of R1 and R2, and 3) a relatively rare “hypersensitive” profile in which R1 and R2 responding was completely suppressed during initial punishment sessions, but became highly discriminated in later sessions. Crucially, multivariate analysis of behaviour in the task provided critical insight into reward motivation, aversive motivation, and punishment association learning, and the key conclusion was that punishment-sensitive vs. -insensitive phenotypes did not differ in their reward or aversive motivations, and instead primarily differed in their ability to learn Action-Punisher associations. Supporting the translational relevance of these behavioural phenotypes, these findings have been replicated in humans (Jean-Richard-dit-Bressel et al., 2021, 2023, 2024) and shown to be stable across a 6-month interval (Zeng et al., 2025).

A prevailing question is how brain processes contribute to these behavioural differences. One brain region that is strongly implicated in punishment sensitivity is the prefrontal cortex (PFC). Damage to the ventromedial PFC (vmPFC) in humans has been associated with insensitivity and persistence in detrimental behaviours (Bechara et al., 2000; Clark et al., 2008; Fellows & Farah, 2003). Similarly, rodent studies demonstrate that punishment avoidance can be impaired through inactivation of the medial PFC structures, such as medial orbitofrontal (mOFC), infralimbic (IL), or prelimbic (PL) cortex (Chen et al., 2013; Ishikawa et al., 2020; Ma et al., 2020; Resstel et al., 2008). Of note, Ma et al. (2020) used conditioned punishment and showed lesions of rat mOFC prior to punishment impaired acquisition of punishment avoidance without affecting fear to the CS+, paralleling the punishment-insensitive phenotype described in Jean-Richard-dit-Bressel et al. (2019). Lateral portions of PFC, such as lateral orbitofrontal (lOFC) and anterior insular (aIC) cortex have also been implicated in punishment avoidance (Y. Chen et al., 2022; Ishikawa et al., 2020; Jean-Richard-dit-Bressel & McNally, 2016). Together, these findings implicate the PFC subregions in punishment-related learning and behaviour, but their relation to individual differences in punishment sensitivity remains poorly understood.

This study aimed to further explore roles for PFC in punishment sensitivity using the conditioned punishment task. Given previous findings that mOFC lesions induced punishment insensitivity (Ma et al., 2020), we sought to examine whether upregulating mOFC activity via chemogenetic activation could enhance punishment-sensitivity. Histology and other analyses indicated our chemogenetic manipulation was ineffective (see **Supplementary materials**). Therefore, the current study focused on examining relationships between phenotypic behaviour and resting-state transcriptional activity (cFos expression) across frontal regions.

## Results

### Overall behaviour in the conditioned punishment task

Long-Evans rats (*N*=70 [35 female]) underwent the conditioned punishment task (**Figure 1A**). First, animals were trained to press two levers (R1, R2) for food on a VI30s schedule (pre-punishment). In later conditioned punishment sessions, animals could still press the two continuously presented levers for food, but presses on each lever could yield cues on a separate VI30s schedule. R1 yielded a CS+ cue that co-terminated with aversive footshock, whereas R2 yielded a distinct CS-cue with no further consequence.

**Figure 1.**
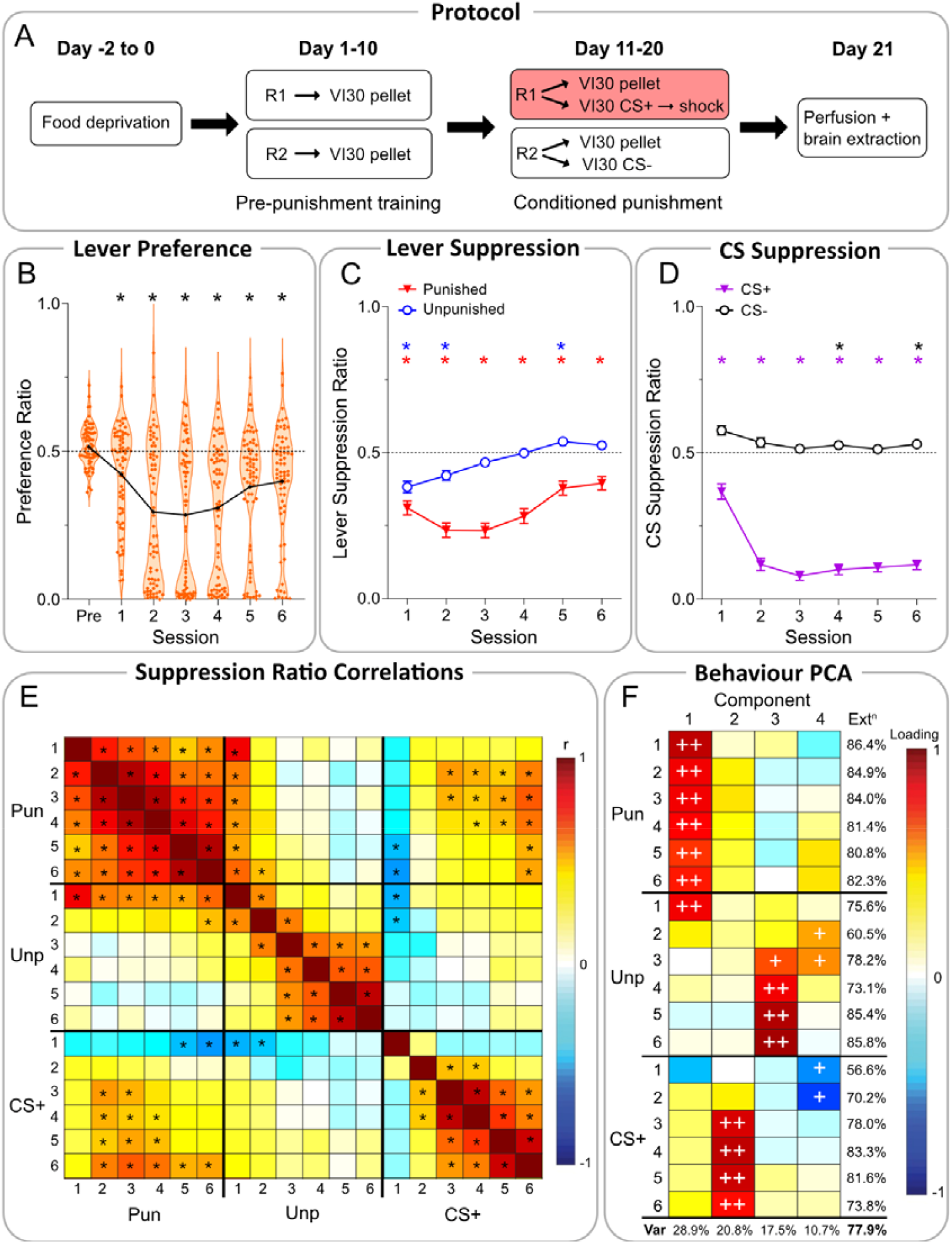
Conditioned punishment task behaviour. **[A]** Task timeline. **[B]** Violin plot of lever preference ratios across sessions (black line = overall mean; orange dots = individuals). \**p*<.01 one-sample t-test against 0.5 (no lever preference). **[C]** Mean ± SEM lever suppression ratios per lever (punished R1, unpunished R2). \**p*<.01 one-sample t-test against 0.5 (no change relative to training). **[D]** Mean ± SEM CS suppression ratios per CS averaged across both levers. \**p*<.01 one-sample t-test against 0.5 (no change relative to non-CS lever-press rates). **[E]** Correlations between punished lever suppression, unpunished lever suppression, and CS+ suppression across conditioned punishment sessions (1-6). \**p*<.05 (FWER-corrected). **[F]** Principal component analysis (PCA) of suppression measures across conditioned punishment sessions (1-6). Measures that loaded substantially (>0.7) or moderately (>0.5) onto each component are indicated with ++ and +, respectively. Bottom row indicates proportion of total variance (Var) accounted for by components. Last column indicates total variance of each measure accounted for by components (extraction).

Punishment avoidance was assessed by calculating a preference ratio between punished R1 vs. unpunished R2 response rates during non-CS periods only (inter-trial interval [ITI]) (**Figure 1B**). A ratio of 0.5 indicated no preference between levers, and a ratio <0.5 indicated a preference for the unpunished lever (i.e., punishment avoidance). Before conditioned punishment, animals showed mild preference for the to-be-punished lever (|*t*_(69)_|=2.320, *p*=.023). With the introduction of the punishment contingency, animals generally learned to avoid the punished lever relative to unpunished lever (Session[linear]: *F*_(1,69)_=12.85, *p*<.001, *η*^*2*^=.309), with preference being significantly <0.5 in all conditioned punishment sessions (|*t*_(69)_|≥3.912, *p*<.001).

Analysis of suppression per lever (relative to pre-punishment lever-press rates) revealed animals significantly suppressed punished (|*t*_(69)_|≥4.543, *p*<.001) but not unpunished responding (|*t*_(69)_|≥ .143, *p*≤.443) across all days. Response suppression during CS presentation was assessed using suppression ratios. Lever-pressing during CS was compared against ITI periods, such that a ratio <0.5 indicated decreased responding during CS presentations relative to ITI (i.e., behavioural suppression). Across all sessions, animals significantly suppressed responding (ratio <0.5) during the aversive CS+ (|*t*_(69)_|≥5.091, *p*<.001) but not CS- (|*t*_(69)_|≥.914, *p*≤.364). Despite exhibiting strong and robust CS+ aversion, animals were highly variable in their punishment avoidance and followed a bimodal distribution, similar to previous studies (**Figure 1B**).

### Underlying components of behaviour within conditioned punishment

We next examined correlations across punished, unpunished, and CS+ suppression ratios to better understand relationships between punishment avoidance, unpunished reward-seeking, and fear learning across sessions (**Figure 1E**). Majority of significant correlations were within-measure (across sessions) and not between-measures, suggesting the processes dictating underlying punishment avoidance, reward-seeking, and fear are largely dissociated from one another. However, there were between-measure correlations, indicative of some interactions between punishment, reward, and fear in driving overall behaviour.

To further understand this, we applied principal components analysis (PCA) to extract orthogonal components of variance across measures to identify latent processes. Four components extracted ∼78% of all variance across subjects (**Figure 1F**). The first component captured variability in punished R1 avoidance, which accounted for 28.9% of total variance. The second component captured variability in eventual CS+ fear (20.8% total variance). The third component captured variance in eventual unpunished R2 responding (17.5% total variance). Finally, the fourth component (10.7% of total variance) had minor positive loadings from mid-punishment R2 responding and negative loadings from initial CS+ suppression, attributable to contextual fear (see Hypersensitive cluster described below).

So, identical to earlier studies, behaviour in the conditioned punishment task was decomposable into independent psychological processes of punishment avoidance (Component 1), Pavlovian aversion (Component 2), unpunished reward-seeking (Component 3), and early contextual fear (Component 4).

### Punishment sensitivity phenotypes

We next characterised key profiles of behaviour in the task using K-means clustering. Punished, unpunished, and CS+ suppression ratios across sessions were used as inputs. Silhouette values were obtained for a range of cluster solutions (*k* = 2-7) (**Figure S1A**). 2-cluster and 4-cluster solutions had unanimously positive values, indicating all subjects were well-accounted for by these solutions. The 2-cluster solution broadly identified punishment-sensitive (*n*=33) vs. insensitive (*n*=37) phenotypes (**Figure S1B-C**). The 4-cluster solution further split these punishment-sensitive vs. insensitive clusters to reveal qualitatively distinct, psychologically-definable sub-clusters, including the previously-described hypersensitive cluster (**Figure 2**). Given the additional insights offered by the 4-cluster solution, we use this solution to identify behavioural phenotypes in this study. Based on behavioural profiles, we labelled these clusters Sensitive (*n*=23), Hypersensitive (*n*=10), Insensitive (*n*=27), and Generalised (*n*=10).

**Figure 2.**
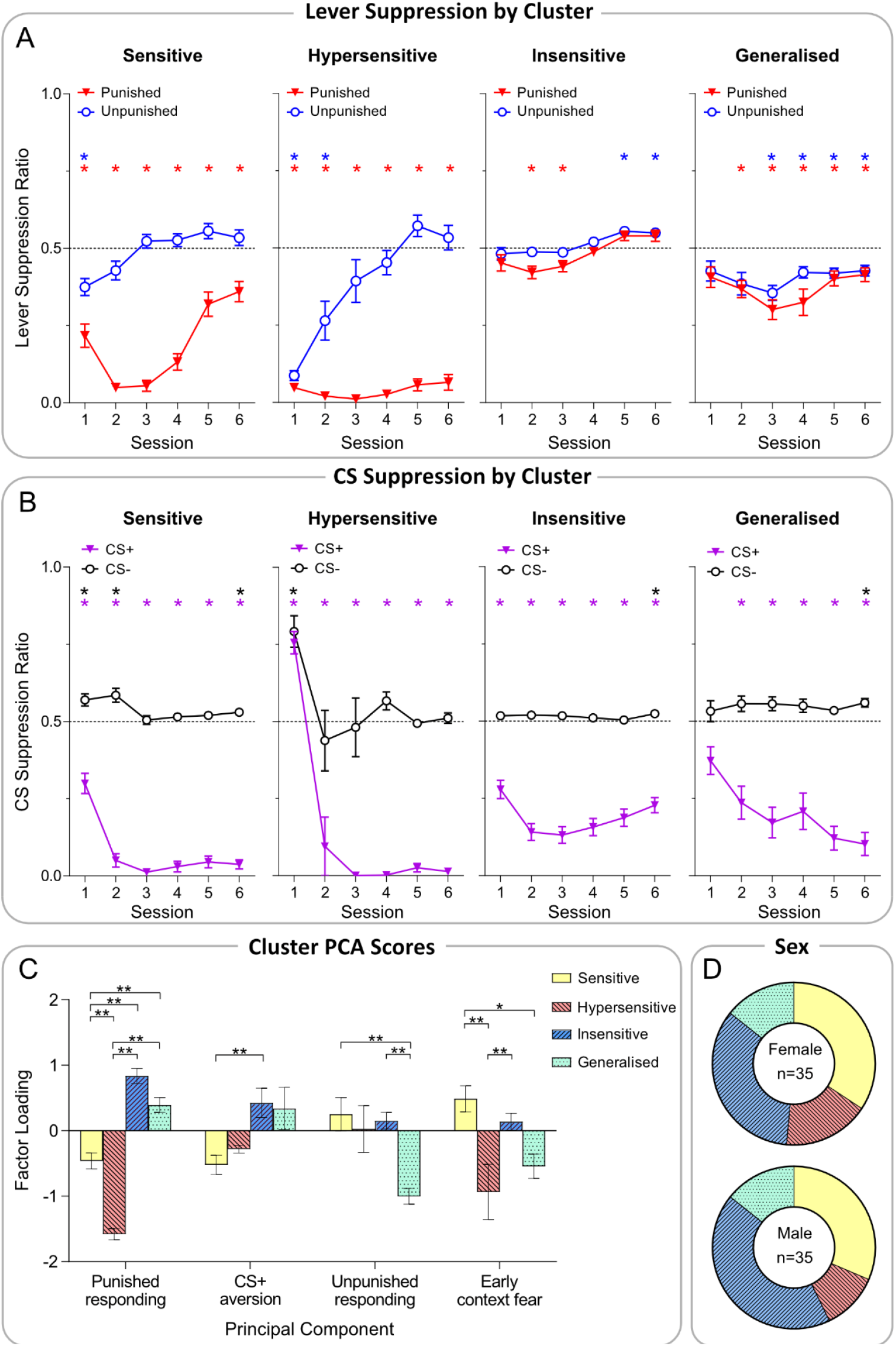
Cluster differences in behavioural performance. **[A]** Mean ± SEM lever suppression ratios per lever (punished R1, unpunished R2), per cluster. \**p*<.01 one-sample t-test against 0.5 (no change relative to training). **[B]** Mean ± SEM CS suppression ratios per CS averaged across both levers. \**p*<.01 one-sample t-test against 0.5 (no change relative to non-CS lever-press rates). **[C]** Mean ± SEM per-individual factor scores (extracted from principal component analysis), per cluster. Significant pairwise comparisons (Tukey-corrected) indicated by \**p*<.05 and \*\**p*<.01. **[D]** Breakdown of cluster assignment across males and females.

Clusters were mainly distinguished by how they suppressed lever pressing across punishment sessions (Cluster*Lever: *F*_(3,66)_=55.66, *p*<.001, *η*^*2*^=.717; Cluster*Lever*Session: *F*_(3,66)_=13.91, *p*<.001, *η*^*2*^=.387) (**Figure 2A**). Sensitives learned to selectively avoid punished relative to unpunished lever (Lever: *F*_(1,22)_=154.7, *p*<.001, *η*^*2*^=.876), exhibiting discriminated lever suppression across all sessions relative to baseline responding (Punished: |*t*_(22)_|≥4.223, *p*<.001; Unpunished: |*t*_(22)_|≥1.045, *p*≤.307). Interestingly, Sensitives exhibited a quadratic trend in punished responding across sessions (Lever*Session: *F*_(1,22)_=61.40, *p*<.001, *η*^*2*^ =.736), suggesting habituation to punishment in later sessions, although punished responding remained significantly suppressed. Hypersensitives indiscriminately suppressed responding in early sessions, before showing strong discrimination (Lever: *F*_(1,9)_=154.7, *p*<.001, *η*^*2*^=.914). Unpunished responding was initially suppressed relative to baseline rates (Session 1-2: |*t*_(9)_|≥3.731, *p*≤.005) and later returned to baseline (Session 3-6: |*t*_(9)_|≥.845, *p*≤.420), while punished responding was suppressed across all sessions (|*t*_(9)_|≥16.95, *p*<.001).

By contrast, Insensitive (Lever: *F*_(1,26)_=6.723, *p*=.015, *η*^*2*^=.205) and Generalised (Lever: *F*_(1,9)_=4.288, *p*=.068, *η*^*2*^=.323) clusters largely failed to discriminate between levers. Insensitive and Generalised clusters were distinguished by whether they suppressed responding during punishment sessions relative to pre-punishment rates. Insensitives broadly maintained response rates on both levers across punishment, and failed to suppress responding compared to baseline (Punished: |*t*_(26)_|=1.508, *p*=.144; Unpunished: |*t*_(26)_|=1.315, *p*=.100). By contrast, the Generalised cluster exhibited generalised response suppression (hence their label) relative to pre-punishment rates (Punished: |*t*_(9)_|=6.775, *p*<.001; Unpunished: |*t*_(9)_|=5.864, *p*<.001), attributable to contextual fear or undiscriminated punishment learning (O’Brien & Frick, 1996; Seligman, 1972).

Although all clusters showed strong selective fear to CS+ vs. CS- (**Figure 2B**), there were significant cluster differences in CS-elicited suppression (Cluster*CS: *F*_(3,66)_=8.101, *p*<.001, η^2^ =.269; Cluster*CS*Session: *F*_(3,66)_=10.34, *p*<.001, *η*^2^ =.457). Further analyses revealed clusters differed in CS+ (Cluster: *F*_(3,66)_=8.042, *p*<.001, *η*^*2*^=.268) but not CS-suppression (Cluster: *F*_(3,66)_=2.056, *p*=.115, *η*^*2*^=.085). All clusters, except Hypersensitives, exhibited significantly greater suppression to CS+ than CS-across all sessions (Sensitive: |*t*_(22)_|≥7.602, *p*<.001; Insensitive: |*t*_(26)_|≥7.563, *p*<.001; Generalised: |*t*_(9)_|≥4.926, *p*<.001). Hypersensitives intially showed similar suppression to both CSs (Session 1-2: |*t*_(22)_|≤2.057, *p*≥.07) before later distinguishing and suppressing significantly more to CS+ than CS- (Session 3-6: |*t*_(22)_|≥5.044, *p*<.001). This effect is likely driven by low initial press-rates seen in Hypersensitives – low responding (due to contextual fear) impairs any clear, dissociable effects between CSs that would otherwise be observed in individuals with higher response rates. Significant phenotypic differences in CS suppression may also be caused by Insensitives exhibiting a quadratic trend in CS+ suppression (Session[quadratic]: *F*_(1,26)_=15.70, *p*<.001, *η*^*2*^=.377), suggesting possible habituation to shock in later sessions.

Differences in behavioural patterns between phenotypes were further explored using per-individual PCA factor scores (**Figure 2C**). Unsurprisingly, clusters differed significantly across factors (Cluster: *F*_(3,66)_=26.01, *p*<.001, *η*^*2*^=.542; Cluster*Factor: *F*_(3,66)_=21.00, *p*<.001, *η*^*2*^=.488). Follow-up pairwise comparisons revealed significant differences in punished responding (Factor 1) between almost every pair, further confirming our cluster analysis. Interestingly, Sensitives and Insensitives differed significantly in their CS+ aversion loadings (Factor 2) which is likely caused by shock habituation in Insensitive animals. Clusters also significantly differed on unpunished responding (Factor 3) and early contextual fear (Factor 4), largely resulting from indiscriminate response suppression seen in both Generalised and Hypersensitive animals.

Although there were slight trends towards females being over-represented in Hypersensitives and males being over-represented in Insensitives, sex was not a significant factor on cluster distributions (**Figure 2D, S1D**; *Χ*^2^_(3)_=.777, *p*=.855).

In summary, we broadly replicate previous observations in a larger cohort of animals. We show that conditioned punishment behaviour is highly variable, in large part due to phenotypic variations in punished response suppression, and to a lesser extent generalised unpunished response suppression. We also observe the same 4 components of behaviour – punishment, Pavlovian fear, reward-seeking, and contextual fear – seen previously (Jean-Richard-dit-Bressel et al., 2019).

### PFC c-Fos across punishment phenotypes

We next examined PFC signatures for behavioural differences via transcriptional activity marker c-Fos. On the day following the final conditioned punishment session (**Figure 1A**), animals’ brains were formaldehyde-fixed, sectioned coronally through PFC, and stained for c-Fos via immunohistochemistry. Scanned sections were registered to the Waxholm-Space Rat Brain Atlas (Kleven et al., 2023) (**Figure 3A**) and c-Fos+ cells were quantified using the QUINT workflow (Yates et al., 2019). Three animals were excluded from analyses due to poor c-Fos registration (<50 c-Fos cells per hemisphere), leaving *N*=67.

**Figure 3.**
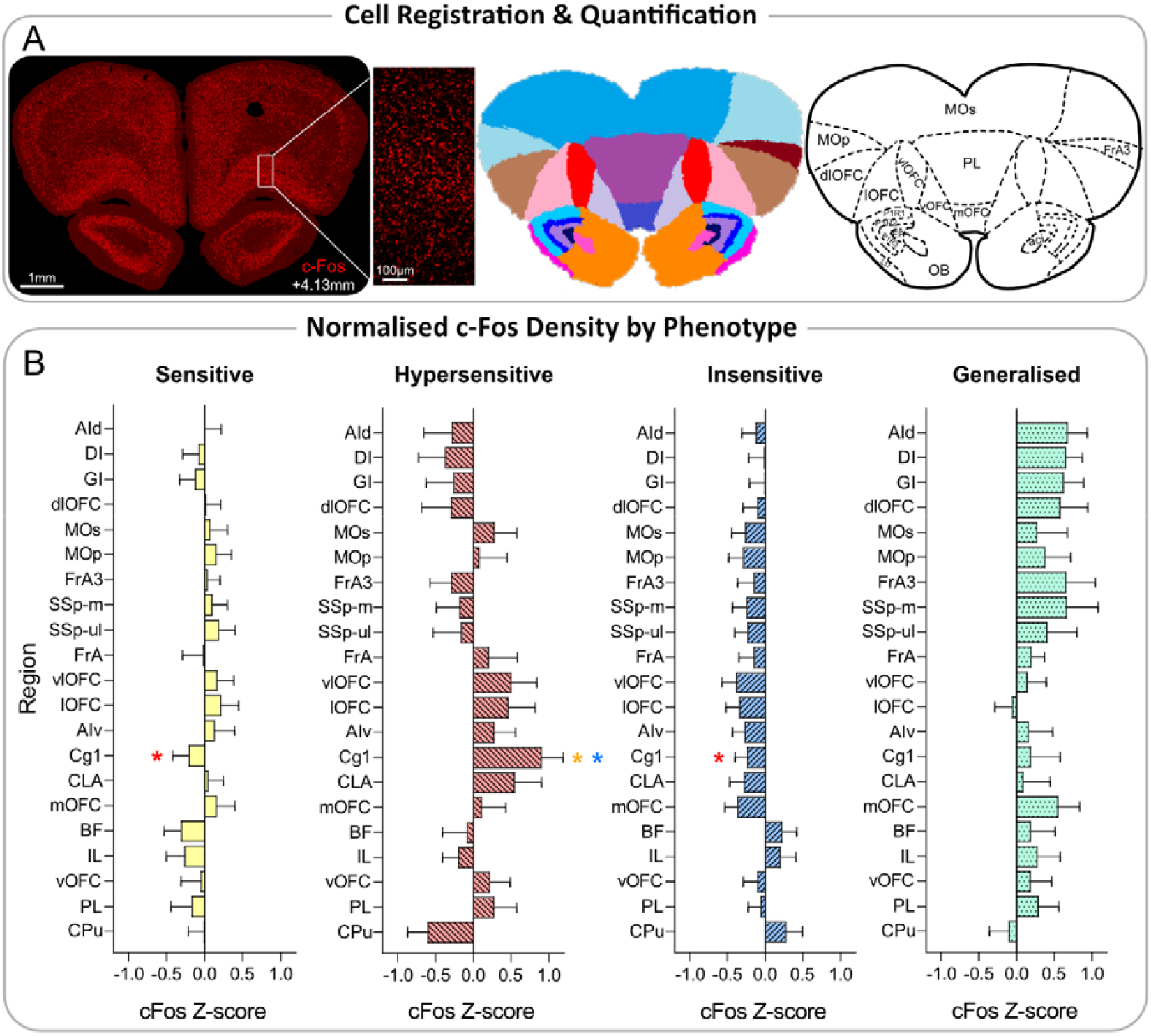
Medial PFC c-Fos density. **[A]** Registration and quantification of c-Fos+ cells via Waxholm-Space atlas. Left: representative photograph of c-Fos+ cells with close-up image (+4.13mm from the decussation of the anterior commissure). Scale = 1mm and 100μmm respectively. Middle: segmentation of regions, output from QuPath InstanSeg extension. Right: segmentation with region labels. **[B]** Mean ± SEM normalised c-Fos density z-scores per cluster. \**p*<.01 significant pairwise comparisons per region between groups. *colour corresponds with cluster (red = different to hypersensitives, yellow = different to sensitives, blue = different to insensitives).

The first question was whether punishment phenotypes showed differential activity in any particular PFC region. ANOVAs on c-Fos density per region only indicated a significant effect of cluster in anterior cingulate cortex (Cg1) (**Figure 3B**; *F*_(3,63)_=4.195, *p*=.009, *η*^*2*^=.166). Follow-up pairwise comparisons of Cg1 activity (Tukey-corrected) revealed Hypersensitives had significantly higher Cg1 activity than Insensitives (*p*=.009) and Sensitives (*p*=.014). Region-specific ANOVAs did not reveal any other significant cluster differences, although differences in mOFC (*F*_(3,63)_=2.481, *p*=.069, *η*^*2*^=.106) and ventrolateral orbitofrontal cortex (vlOFC) (*F*_(3,63)_=2.567, *p*=.062, *η*^*2*^=.109) approached significance.

So, despite stark behavioural differences, there were limited differences in PFC c-Fos when analysed by region. Additionally, this broad analysis fails to provide insight into the specific behavioural aspects driving differences observed in Cg1 c-Fos. However, contemporary accounts of brain function emphasise recruitment of large-scale, overlapping brain networks, over isolated activity patterns within individual regions, in understanding how the brain shapes behaviour (Rubinov & Sporns, 2010; Sporns et al., 2007; Terstege & Epp, 2023). We therefore applied multivariate approaches to extract PFC networks and examine how activity in these networks related to phenotypic behaviour.

PCA was employed to extract components of covariance in c-Fos across frontal regions to identify co-activation networks (**Figure 4A**). Ten animals were excluded from this analysis (*N*=57) due to missing data for 1 or more regions. Regions loaded strongly onto 3 main components (i.e., networks), where activity of one region could predict activity of other regions within the same network. Network 1 (N1) accounted for 32.8% of total variance in frontal activity and consisted of dorsal agranular insular cortex (AId), dysgranular insular cortex (DI), granular insular cortex (GI), dorsolateral orbitofrontal cortex (dlOFC), secondary motor cortex (MOs), primary motor cortex (MOp), frontal association area 3 (FrA3), primary somatosensory areas (SSp-m, SSp-ul), and frontal association cortex (FrA). Network 2 (N2) accounted for 21.5% of variance and consisted of ventrolateral orbitofrontal cortex (vlOFC), lateral orbitofrontal cortex (lOFC), ventral agranular insular cortex (AIv), cingulate area 1 (Cg1), and the claustrum (CLA). Finally, Network 3 (N3) accounted for 16.4% of variance and consisted of medial orbitofrontal cortex (mOFC), ventral orbitofrontal cortex (vOFC), basal forebrain region (BF), infralimbic cortex (IL), prelimbic cortex (PL), and caudate putamen (CPu). Additional hierarchical clustering analysis revealed the same 3 clusters when cut at half height, further supporting the classification of these networks (**Figure 4B**). N1 mostly consisted of motor and somatosensory areas, while N2 and N3 predominantly contained ventrolateral and medial PFC structures respectively (**Figure 4C**).

**Figure 4.**
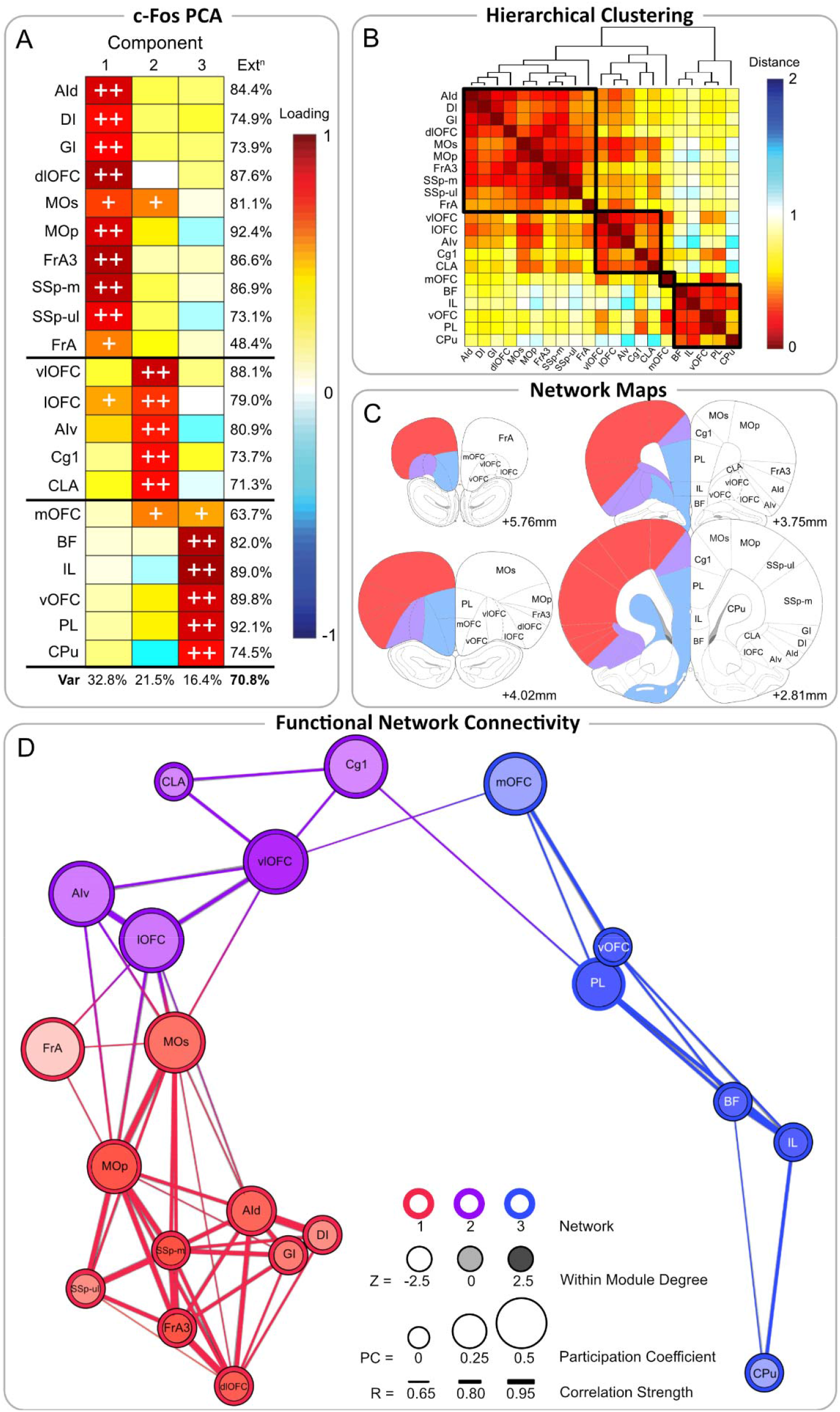
Functional connectivity network analyses. **[A]** Principal component analysis of regional c-Fos density. Regions that loaded substantially (>0.7) or moderately (>0.5) onto each component are indicated with ++ and +, respectively. Bottom row indicates proportion of total variance (Var) accounted for by components. Last column indicates variance of each measure accounted for by components (extraction). **[B]** Hierarchical clustering of the distance matrix (1-*r*) based on Pearson’s correlational coefficient *r* between every pair of regions. Dendrogram was cut at half height to identify network clusters (outlined in black squares). **[C]** Representative coronal diagrams of network assignment, based on Waxholm-Space Atlas. Red colour = Network 1, purple colour = Network 2, blue colour = Network 3. Diagrams are +5.76mm, +4.02mm, +3.75mm, and +2.81mm from the decussation of the anterior commissure (top to bottom, left to right, respectively). **[D]** Functional connectivity graph, thresholded to *r*≥0.65. Circles depict nodes/brain regions, with connecting lines being edges/correlational strength between regions. Circle colour represents network assignment (red = Network 1, purple = Network 2, blue = Network 3). Circle colour saturation represents within-module degree Z-score (pale colour = lowest, dark colour = highest). Circle size represents participation coefficient scores (smallest size = lowest, biggest size = highest). Line thickness represents correlational strength (thinner = weaker, thicker = stronger).

We also applied a graph theory approach to further characterise these networks and identify how each region participates within and across these networks (**Figure 4D**). To measure inter- and intra-network connectivity, we calculated participation coefficient (PC; the extent to which regions are connected to regions in other networks, relative to those in the same network), and within-module degree Z-score (WMDz; the extent to which each region is connected to regions within the same network) respectively (Guimerà & Nunes Amaral, 2005; Kimbrough et al., 2020). N2 had the highest inter-module connectivity with 4 regions having a high PC (vlOFC [.47], lOFC [.47], AIv [.50], Cg1 [.44]), while N1 had 2 regions (MOs [.42], MOp [.30]), and N3 had 1 region (mOFC [.42]). N1 had the highest intra-module connectivity with 3 regions having a high WMDz (MOp [.84], FrA3 [.87], SSp-m [.96]), while N2 had 1 region (vlOFC [1.76]), and N3 had 2 regions (vOFC [.70], PL [.72]). Notably, MOp (N1) and vlOFC (N2) had both high PC and WMDz values, indicating the importance of their role both within- and between-networks. Taken altogether, these results suggest N1 and N3 largely interact within their respective networks, while N2 acts as a bridge to relay information between networks.

Critical hub regions were identified via a normalised between-ness centrality score (BC), which calculates how often a region lies on the shortest path between all pairs of regions (Freeman, 1977, 1978; Kintali, 2008). Regions with higher BC scores are considered to have an increased capacity to facilitate activity between regions and are crucial to the overall structure of the network (Rubinov & Sporns, 2010; Terstege & Epp, 2023). The top 3 regions with the highest BC scores were the vlOFC (BC=.51), MOp (BC=.29), and mOFC (BC=.34). These regions each have high PC scores and are members of separate networks, suggesting vlOFC, MOp, and mOFC are critical hub regions for inter-network communication.

### Network activity predicts behaviours associated with punishment and contextual fear

Next, we explored relationships between network activity and phenotypic behaviour by assessing cluster differences in network activity scores. There were no significant overall cluster differences in N1 (*F*_(3,56)_=1.836, *p*=.151, *η*^*2*^=.090) nor N3 (*F*_(3,56)_=.697, *p*=.564, *η*^*2*^=.035) activity. However, clusters significantly differed in N2 activity (*F*_(3,56)_=4.067, *p*=.011, *η*^*2*^=.179), with Hypersensitives having significantly higher N2 scores than Insensitives (**Figure 5A**; *p*=.007), suggesting N2 may be implicated in punishment avoidance and/or behavioural suppression.

**Figure 5.**
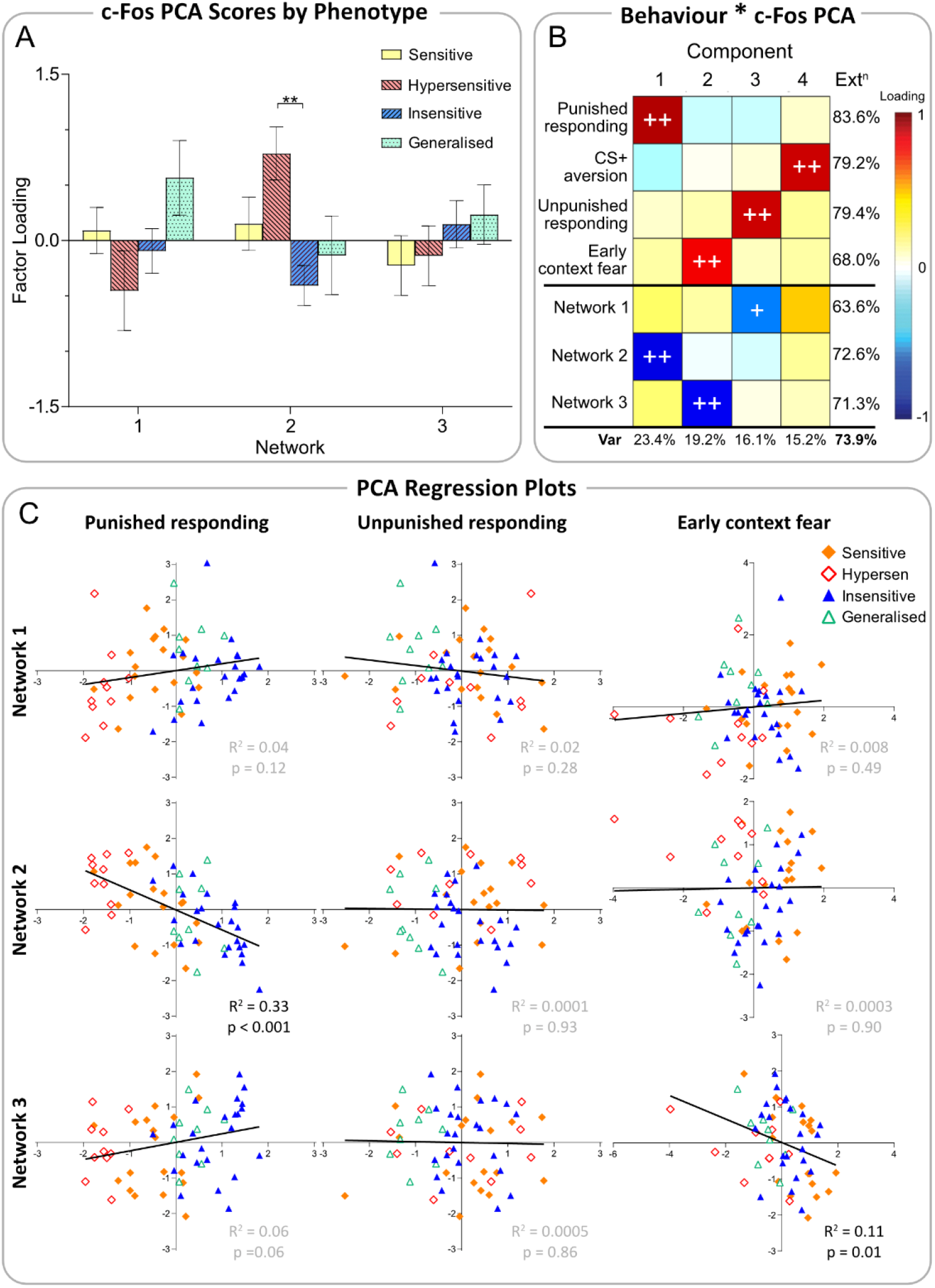
Relationship between network activity and behaviour. **[A]** Mean ± SEM per-individual c-Fos factor scores (extracted from principal component analysis), per cluster. Significant pairwise comparisons (Tukey-corrected) indicated by \*\**p*<.01. **[B]** Principal component analysis of per-individual factor scores (extracted from behavioural and c-Fos principal component analyses). Factors that loaded substantially (>0.7) or moderately (>0.5) onto each component are indicated with ++ and +, respectively. Bottom row indicates proportion of total variance (Var) accounted for by components. Last column indicates variance of each factor accounted for by components (extraction). **[C]** Linear regression analyses with line of best fit between per-individual behavioural factor scores and per-individual c-Fos factor scores (extracted from respective principal component analyses). Each point represents a separate individual and colour/symbol indicates cluster assignment. Respective *R*^*2*^ and *p*-values located in the bottom right for each graph (significant relationships in black text, non-significant relationships in grey).

To further determine how network activity relates to behavioural measures, we conducted PCA on behavioural component scores and c-Fos network scores described previously (**Figure 5B**). This analysis revealed that component scores for punished responding were strongly and inversely related to N2 activity, indicating that animals with increased N2 activity exhibited greater punished response suppression (i.e., Hypersensitives). This corroborates our previous finding of significant cluster differences in N2 activity. Interestingly, scores for early contextual fear were strongly and inversely related to N3 activity, despite the absence of significant cluster differences in N3 activity. There was also a relatively weak relationship between scores for unpunished responding with N1 activity, and CS+ aversion did not meaningfully load onto frontal network activity. This suggests that distinct components of behaviour are related to the activity of separately-identified frontal networks.

We directly examined these relationships in a pairwise manner using linear regression (**Figure 5C**). In agreement with PCA results, increased N2 activity was associated with decreased punished responding (*R*^*2*^=.33, *p*<.001), while increased N3 activity was associated with early contextual fear (i.e., generalised response suppression) (*R*^*2*^=.11, *p*=.01). N1 activity was not significantly associated with any behavioural component (punished responding [*R*^*2*^=.04, *p*=.12], early contextual fear [*R*^*2*^=.0008, *p*=.49], unpunished responding [*R*^*2*^=.02, *p*=.28]). This is relatively unsurprising, given N1 did not load strongly with any behavioural measure.

## Discussion

The present study sought to investigate whether variance in punishment-related behaviour was reflected in distinct brain activity patterns in the prefrontal cortex. To do this, we leveraged the conditioned punishment task, and broadly replicated all key observations from the original individual differences study (Jean-Richard-dit-Bressel et al., 2019): 1) the bimodal distribution of punishment avoidance, 2) punishment-dissociated phenotypes, and 3) the latent 4-component structure of psychological processes driving behaviour in the task. Analysis of prefrontal c-Fos revealed few differences in c-Fos expression across phenotypes when examined on a per-region basis, but further analyses revealed differential engagement of 3 PFC networks. Specifically, activity in a network primarily composed of ventrolateral PFC regions (Network 2) strongly predicted punishment avoidance, while activity in a network of predominantly medial PFC regions (Network 3) predicted initial generalised response suppression. We elaborate on these findings below. The replication of all key conditioned punishment behavioural findings in a new larger cohort of animals indicates the robustness of original findings, and the utility of this task for investigating individual differences in punishment and related behaviours. One notable difference between study findings is that the original study identified 3 behavioural profiles, whereas the current study identifies 4. The current study’s Sensitive, Generalised, and Hypersensitive phenotypes conform to “punishment-sensitive”, “punishment-insensitive”, and “hyper-sensitive” phenotypes in the previous study. However, the current study highlights an additional behavioural profile, termed the Insensitive phenotype here. In contrast to this study’s Generalised cluster, who failed to bias responding away from punished lever but showed modest response suppression across punishment, the Insensitive cluster did not suppress punished or unpunished responding across punishment sessions, and showed the least fear to the aversive CS+ by the end of punishment. This profile conforms with traditional notions of insensitivity, i.e., reduced “sensing” of the punisher or increased pain thresholds (Corr, 2004; Engeln & Ahmed, 2024), theoretically resulting in reduced aversion to environmental (CS+) and behavioural (punished response) predictors. Detection of this additional phenotype likely results from the current study’s larger sample size (N=70 vs original N=48), which permits statistical dissociation of the relatively subtle distinction between Insensitive vs. Generalised phenotypes.

The only individual region whose c-Fos activity was significantly different across clusters was the anterior cingulate cortex (Cg1), in which Hypersensitives had significantly higher expression than Insensitives and Sensitives. Activity in Cg1 is associated with acquisition and expression of contextual fear (de Lima et al., 2022) and defensive behaviours (Falconi-Sobrinho et al., 2017, 2021, 2024). This aligns with our findings, as Hypersensitives exhibited the greatest general response suppression, indicative of contextual fear, while Insensitives and Sensitives showed the least generalised suppression. The lack of significant c-Fos differences in other punishment-implicated PFC regions (e.g., OFC, PL, IL [Burgos-Robles et al., 2017; B. T. Chen et al., 2013; Jean-Richard-dit-Bressel & McNally, 2016; Ma et al., 2020; Orsini et al., 2015; Piantadosi et al., 2017; Resstel et al., 2008; Verharen et al., 2019]) is somewhat surprising, but additional network-level analyses of PFC activity implicate these regions in ways that are consistent with existing literature.

Ventrolateral PFC structures like lOFC and aIC are implicated in the expression of punishment avoidance (Y. Chen et al., 2022; Ishikawa et al., 2020; Jean-Richard-dit-Bressel & McNally, 2016). Fittingly, activity of the ventrolateral-dominant network (N2) strongly predicted punishment avoidance in the current study. Conversely, medial PFC structures (e.g., PL, IL, medial OFC) are less reliably associated with punishment-related behaviours (see Jean-Richard-Dit-Bressel et al., 2018). Inactivations of PL or IL have been shown to impair behavioural suppression under certain punishment conditions (Broomer & Bouton, 2024; B. T. Chen et al., 2013; Resstel et al., 2008), but also in Pavlovian fear paradigms (Burgos-Robles et al., 2017; Giustino & Maren, 2015; Limpens et al., 2015), suggesting a broader role in aversion sensitivity. Pertinently, medial OFC lesions prior to conditioned punishment specifically impaired punishment suppression but not fear-elicited suppression (Ma et al., 2020), indicating medial PFC networks do not mediate general aversive learning. Instead, these regions may be critical for resolving whether aversive events should be attributed to one’s actions over environmental stimuli. Correspondingly, activity in the medial PFC-dominant network (N3) was most strongly related to the initial partitioning of punishment vs. contextual fear. Finally, N1 did not strongly relate to phenotypic differences in behaviour. N1 was most notably composed of motor and somatosensory cortical areas, which are not typically considered part of PFC.

Contrastingly, variance in CS+ aversion and unpunished responding were not (or only weakly) accounted for by PFC network engagement. This might indicate the greater importance of subcortical circuitry in fear and reward-seeking behaviour (e.g., limbic and basal ganglia circuits, respectively). It is well established that the amygdala is a key limbic structure mediating fear acquisition and expression (Davis, 1992; Grewe et al., 2017; Maren & Quirk, 2004; Sah et al., 2003; Sengupta et al., 2018), and amygdala connectivity to mPFC is implicated in this role (Burgos-Robles et al., 2017; Giustino & Maren, 2015). Similarly, the striatum is a key component of basal ganglia circuitry known for its role in reward-related learning and behaviour (Balleine et al., 2007; Cox & Witten, 2019), with frontal cortex input to striatum being strongly implicated in these functions. It stands to reason that variance in fear and unpunished reward-seeking may be specifically captured by activity of PFC neurons connected with the amygdala and striatum, respectively, but isolation of these specific sub-populations was not possible in the current study. Future work could build upon the findings of the current study using circuit-based approaches to explore these possibilities.

Together, these results suggest punishment sensitivity is not attributable to activity in any single PFC region, but may instead be attributed to differences in patterns of activity across neural networks. This importance of network-level recruitment over individual regions in punishment may explain the lack of reliable effects on punishment avoidance when conducting region-specific manipulations of PFC. Many studies report PFC inactivations increase punished responding (Burgos-Robles et al., 2017; B. T. Chen et al., 2013; Clark et al., 2008; Ishikawa et al., 2020; Jean-Richard-dit-Bressel & McNally, 2016; Ma et al., 2020; Resstel et al., 2008), while others report decreased punished responding (Ishikawa et al., 2020; Liley et al., 2019; Orsini et al., 2015), or no effects (Jean-Richard-dit-Bressel & McNally, 2016; Pelloux et al., 2013). This mixed evidence has been reasonably attributed to differences in task demands (Bouton & Broomer, 2023; Jean-Richard-Dit-Bressel et al., 2018). However, the current findings suggest a more holistic network-level manipulation may be more conceptually sound than individual region-level manipulations. Network-level engagement was the more relevant predictor of punishment behaviour, with network analysis indicating some degree of redundancy for any given PFC region in intra- and inter-region activity. So, manipulations of networks over individual regions might produce more reliable effects on punishment learning and behaviour, potentially addressing the highly conflicting literature around roles for PFC in punishment.

It is important to note that this study has several limitations. First, our findings are merely correlational; future research will be necessary to draw causal links between PFC network recruitment and punishment learning and avoidance. Additionally, animals here received ligand injections in the attempt to chemogenetically manipulate PFC activity, which may have confounded our findings. A slew of analyses suggest this is unlikely, as chemogenetic ligands had no effect on behaviour or c-Fos relative to saline (**Supplementary Materials**), but this remains a caveat of the current study. Finally, brains were obtained the day after the final conditioned punishment session, so c-Fos activity reported here was not directly task-related, instead representing resting transcriptional activity. Resting-state activity has been shown to predict task-relevant neural activity (Reineberg et al., 2015; Smith et al., 2009), but future work examining task-relevant PFC activity, including real-time neural dynamics in response to task components (actions, cues, outcomes), would be an important extension of current findings.

In conclusion, individuals vary substantially in their propensity to reduce harmful behaviours. Here, we replicated the bimodal distribution in punishment avoidance and describe a new punishment-insensitive phenotype in the task. Using PCA, hierarchical clustering, and graph theory approaches, we identified distinct PFC co-activity networks that support orthogonal components of behaviour to produce qualitatively distinct punishment phenotypes. Findings suggest a ventrolateral PFC network promotes punishment avoidance, while a medial PFC network arbitrates initial punishment learning from generalised suppression. Future research is needed to explore the causal relationships between network engagement and punishment sensitivity, which may in turn lead to neural intervention targets for maladaptive punishment sensitivity.

## Methods

### Subjects

Subjects were experimentally-naïve Long-Evans rats (N=70 [35 female]) aged 7-8 weeks. Animals were housed in groups of 4 in a temperature and humidity-controlled room, maintained on a 12-hr light/dark cycle. All behavioural training and experiments occurred during the light phase. Animals were food-restricted up to 2 days before commencement of behavioural procedures, and maintained at ∼90% of their free-feeding weight for the remainder of the experiment.

All procedures were approved by the Animal Care and Ethics Committee at UNSW Sydney and conducted in accordance with the National Health and Medical Research Council Code for the Care and Use of Animals for Scientific Purposes in Australia (2013).

### Apparatus

Behavioural procedures were conducted in standard operant conditioning chambers (Med Associates, Inc.) kept within ventilated, sound attenuating cabinets. The magazine dish was flanked by 2 retractable levers, with a keylight above each lever. Grain pellets (45mg) were delivered into the magazine from an external dispenser, and magazine entries were recorded by infrared photobeam sensors. CSs were a 10s 3kHz tone (via wall-mounted speaker) or 5Hz flashing light (via house light and keylights). The grid floor was connected to an external generator that delivered foot-shocks (0.5s, 0.3-0.5mA).

### Behavioural procedures

#### Lever-press training

Rats first received 10 days of lever-press training. Sessions were designed such that animals acquired equal pressing on both levers. On Days 1-2, rats were presented with both levers simultaneously where every lever press produced a pellet (FR1). Levers retracted once 25 presses were registered on a lever, and session terminated once both levers had retracted or 60mins had elapsed (whichever came first). Every subsequent session terminated once 40mins had elapsed.

On Days 3-6, rats were presented with a single lever which alternated every 5mins. Lever-pressing was reinforced on a VI30s schedule, where a lever-press produced a pellet approx. every 30 seconds, on average. On Days 7-9, sessions were intended to remove any bias/preference animals had for either lever. Rats were again presented with both levers simultaneously, and reinforcement occurred on a RI30s schedule such that pellets were more frequently available on the non-preferred lever and less available on the preferred lever. The final session on Day 10 was used to obtain an unbiased measure of pre-punishment lever-press rates (used in lever suppression ratios during data analysis). Here, both levers were available simultaneously and reinforced on a VI30s schedule.

#### Conditioned punishment

Following training, rats received 10 days of conditioned punishment. Sessions were 50mins long and both levers were extended throughout. Lever-presses were reinforced (VI30s schedule) and each response also had the chance to elicit a CS (VI30s schedule). Presses on the punished lever could elicit an aversive CS+ co-terminating with a foot-shock, while presses on the unpunished lever could elicit a neutral CS-which terminated alone. Shock intensity was initially set at either 0.3mA (*n*=23) or 0.4mA (*n*=47) for the first two sessions, but was incrementally increased by 0.1mA (up to 0.5mA) if animals failed to suppress during the CS+ (CS+ suppression ratio >0.4). Identities for punished/unpunished levers (left vs. right) and CS+/− (light vs. tone) were counterbalanced across animals, and remained the same across sessions.

### Surgery and DREADD manipulations

All procedures relating to hM3D are described in **Supplemental Materials**.

### Immunohistochemistry

Rats were anaesthesised with sodium pentobarbital injections (1mL females, 1.5mL males, i.p.) the day after the final conditioned punishment session (Day 21), and transcardially perfused using 4% paraformaldehyde (PFA) in 0.1M phosphate-buffered saline (PBS). Brains were extracted and post-fixed in PFA overnight at −30°C, then stored in 20% sucrose in PBS for 24hrs. Brains were frozen and coronally sectioned at 40µm through the PFC using a cryostat. Sections were stored in 0.1% sodium azide in 0.1M PBS (pH 7.2).

For immunohistochemical analysis, sections were washed in PBS to remove excess sodium azide, and blocked in Triton X-100 solution for 1-2hrs. Sections were incubated overnight in primary antibodies, and washed in PBS before incubation in secondary antibodies (**Supplemental Materials**). Sections were washed again in PBS, then mounted and cover-slipped.

### Image acquisition and cell quantification

Mounted sections were scanned (Axioscan, Zeiss) and imaged via Zeiss ZEN software. The QUINT workflow (Yates et al., 2019) was used to quantify cells co-expressing the DREADD virus and c-Fos protein.

First, to identify the rostral-caudal position of sections in atlas space, accounting for sectioning angle, sections with clear anatomical landmarks were manually registered to the 3D Waxholm Atlas (version 4; RRID: SCR_017124; Kleven et al., 2023) using QuickNII (version 2.2; RRID: SCR_016854; Puchades et al., 2019). Linear adjustments were then made on the remaining sections. To account for tissue distortion, these registrations were then non-linearly adjusted in VisuAlign (version 0.9; RRID: SCR_017978; Puchades et al., 2019).

Separately, the original images were pre-processed in Fiji (Schindelin et al., 2012) prior to cell quantification. To increase the signal-to-noise ratio between labelled cells and background, we used the ‘pseudo flat field correction, gaussian blur, median’ filters available in Fiji. The processed images were then used for cell detection and classification in QuPath (version 0.6.0; Bankhead et al., 2017). After manually removing artifacts and tissue damage, cells were segmented using the QuPath InstanSeg extension (Goldsborough et al., 2024) and classified using the ‘triangle’ automatic thresholding method in Fiji.

For each animal, the atlas registrations generated from VisuAlign were combined with the object segmentations masks from QuPath using the Nutil Quantifier (version 0.8.0; RRID: SCR_017183; Groeneboom et al., 2020). The resulting output was used to calculate cell density per brain region.

### Data analysis

Data was analysed in IBM SPSS Statistics 30, with Type 1 error rate controlled at α=.05. For the behavioural correlation matrix, error rate was FWER-corrected to α=1.63e^-4^. Correlation and PCA heatmaps were created in MatLab. Graph theory network analyses were also performed and visualised in MatLab. Regression plots were analysed and produced in GraphPad Prism. Animals underwent 10 days of conditioned punishment in total, however, behavioural data was only taken from the first 6 days (prior to the second round of ligand injections).

#### Behavioural measures

Behavioural measures were obtained from lever-press rates on punished and unpunished levers during CS+, CS-, and ITI (non-CS) periods. Punishment avoidance was assessed using a lever preference ratio. This measures lever-press rates (LP) on one lever relative to the other lever, and was calculated per session:

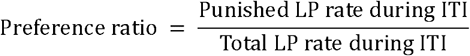

A ratio of 0.5 indicates no preference for either lever, while <0.5 indicates a preference for the unpunished lever (i.e., punishment avoidance).

To assess changes in press rates relative to pre-punishment (last session of lever-press training), lever suppression ratios were calculated per lever per conditioned punishment session:

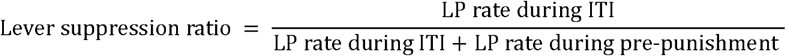

A ratio of 0.5 indicates no change in lever-press rate relative to pre-punishment, while <0.5 indicates reduced lever-pressing (i.e. lever suppression).

To assess Pavlovian fear to CSs, CS suppression ratios were calculated per CS per session:

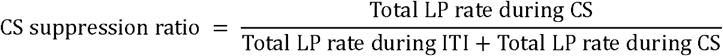

A ratio of 0.5 indicates no behavioural suppression during CS relative to ITI, while <0.5 indicates decreased lever-press rates (i.e. increased suppression) during CS relative to ITI.

#### Punishment sensitivity clusters

We identified punishment sensitivity clusters using a K-means clustering algorithm. Punished, unpunished, and CS+ suppression ratios across sessions were used as input. As described in Results, solutions for k=2-7 were obtained, and the optimal solution was selected based on silhouette values. Solutions for k=2 and k=4 had unanimously positive silhouette values (indicating all animals performed more similarly to others in the same cluster than to animals outside their cluster). The 2-cluster solution identified punishment-sensitive and insensitive clusters. The 4-cluster solution further divided these clusters into sub-types; sensitive and hypersensitive from the sensitive cluster, and generalised and insensitive from the insensitive cluster. We chose the 4-cluster solution because it allowed for deeper insight into individual variance in this task.

#### Task performance

Task performance (lever and CS suppression) across sessions was assessed using repeated-measures ANOVA analyses. Session (Days 1-6), lever (punished vs. unpunished), and CS (CS+ vs CS-) were used as within-subject factors. Cluster assignment (sensitive vs. hypersensitive vs. insensitive vs. generalised) and ligand injections (saline vs. DCZ/CLZ) were used as between-subject factors. Follow-up simple ANOVAs were used to clarify interaction effects. Preference and suppression ratios per session were compared to the null ratio 0.5 using a one-sample t-test.

PCA was used to further analyse relationships between punishment (punished lever suppression), reward (unpunished lever suppression), and fear (CS+ suppression). Results were varimax-rotated to improve interpretability of components. The optimal solution was selected based on total variance accounted for (>70%) and extraction for each measure (>50% for majority of measures). Individual loadings for each component per cluster were extracted and analysed using ANOVA, with punishment sensitivity clusters as the between-subjects factor. Pairwise comparisons were Tukey-corrected.

#### Normalised c-Fos density

c-Fos density for each prefrontal region was obtained as described above. To account for batch differences, densities were adjusted using a linear mixed effects model with cohort as a random effect. Adjusted values were obtained by summing the average intercept across cohorts with the individual residuals. Densities were normalised and Z-scores were calculated per animal per region. ANOVA was used to analyse for cluster differences in Z-scores per region, and pairwise comparisons were Tukey-corrected.

#### Network identification

PCA was conducted to identify co-activating neural regions, measured via c-Fos density. As described above, results were varimax-rotated and the optimal solution was selected based on total variance accounted for (>70%) and extraction for each region (>50% for majority of regions). Hierarchical clustering was used to verify PCA region groupings. Functional distance between regions was computed by calculating (1-*r*) with *r* being Pearson’s correlation coefficient for each pair of regions. “Network” clusters were obtained by cutting the dendrogram at half height, and region order was reorganised based on hierarchical clustering for Z-score and PCA visualisations.

#### Graph theory network analyses

Graph theory is a branch of mathematics that examines relationships in complex networks, and is increasingly being used to identify key brain regions in functional neural connectivity (Kimbrough et al., 2020; Rubinov & Sporns, 2010; Sporns et al., 2007; Terstege & Epp, 2023; Vetere et al., 2017; Wheeler et al., 2013). In graph terms, ‘node’ refers to brain region and ‘edge’ refers to the connections between nodes. We used 3 centrality measures to investigate the importance of each region, both within their respective network and within the entire structure. WMDz measures the extent to which a region is connected to other regions within its network, thus intra-network connectivity, while PC measures the extent to which a region is connected to regions outside its network, thus inter-network connectivity (Guimerà & Nunes Amaral, 2005). Finally, between-ness centrality measures how influential a region is to the overall structure (Freeman, 1977, 1978; Kintali, 2008).

First, we created a correlation matrix using Pearson’s *r* values for every pair of regions. This correlation matrix was used to produce an adjacency matrix, which dictates whether an edge exists between any node pair. A value of ‘1’ was entered for moderate-strong correlations (*r*≥0.65) and ‘0’ for values of r<0.65. Hence, only edges with positive weights ≥0.65 were visualised and analysed. We initially tried to include only strong positive correlations (*r*≥0.75) as has previously been used (Kimbrough et al., 2020), however this excluded FrA from having any edges/connections to other regions. Therefore, we chose 0.65 because that was the highest value we could use while still including all regions. We used a weighted adjacency matrix (calculated by multiplying the adjacency and correlation matrices) for graph visualisation and metric calculation. Multidimensional scaling analysis was conducted on the functional distance between regions (1-*r*). This produced arbitrary Cartesian coordinates, allowing us to plot the relative position of each region such that highly correlated regions would appear closer together.

WMDz and PC measures were calculated using formulae previously defined for graphs with weighted edges (Guimerà & Nunes Amaral, 2005; Kimbrough et al., 2020). For WMDz, we first calculated the within-module degree (*k*_*i*_) for each region (*i*) by summing the weights of all edges between region (*i*) and other regions within the same network (*s*_*i*_). To calculate each region’s within-module degree Z-score, the average within-module degree for that network 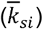 was subtracted from the region’s within-module degree, and divided by the standard deviation for that network (*σ*_*ksi*_). Hence, WMDz is given by

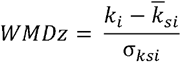

For PC, we first calculated the between-module degree (*k*_*is*_) for each region (*i*) by summing the weights of all edges between region (*i*) and regions in network (*s*). This was then divided by the total degree (*k*_*i*_), defined as the summed weight of all edges between region (*i*) and all other regions, and squared. This process was repeated for each of the 3 networks (*s*=1 to *s*=3), summed, and subtracted from 1. Hence, the PC for each region is given by

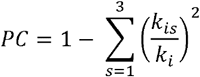

Nodes were considered to have high intra-network connectivity if WMDz ≥0.70, and high inter-network connectivity if PC ≥0.30 as has previously been used in Fos-related analyses (Kimbrough et al., 2020). In the current data, WMDz values ranged from −2.26 to 1.78, with all regions (except vlOFC) scoring <1.00. PC values ranged from 0 to 0.50, with 11/21 regions scoring 0.

Between-ness centrality (BC) is a popular measure of node importance, and refers to how often a node lies on the shortest path between any pair of nodes (Freeman, 1977, 1978; Kintali, 2008). Nodes with higher BC scores are considered more crucial to the graph structure because they facilitate communication between more nodes – i.e., if such nodes were deleted from the graph, then many of the remaining nodes would need to communicate via a slower and less efficient path. In contrast, deleting nodes with lower BC scores would have a lesser effect on connections between remaining nodes. Centrality scores for each region (*v*) were calculated using an inbuilt graph centrality function in MatLab. First, the shortest paths between any pair of nodes (*s,t*) were found, accounting for edge weights such that paths between highly correlated regions (i.e., larger edge weights) were preferred over paths between less correlated regions (i.e., smaller edge weights). The total number of shortest paths passing through the region (*λ*_*st*_*(v)*) is then calculated as a fraction of the total number of shortest paths (*λ*_*st*_) for that node pair. This process was repeated for every possible node pair and summed, producing the centrality score for that region. Finally, centrality scores were divided by the total number of node pairs to normalise values. Hence, BC is given by

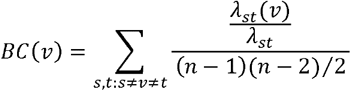

For the current study, we only reported the top 3 regions with the highest BC values. BC values ranged from 0 to 0.52, with 9/21 regions scoring 0.

#### Behaviour x network analyses

To assess whether network activity differed across punishment sensitivity clusters, individual loadings for each c-Fos PCA component (i.e., network) were extracted and analysed using ANOVA, with punishment sensitivity clusters as the between-subjects factor. Follow-up pairwise comparisons were Tukey-corrected. However, this did not yield much result. To further examine any underlying relationships between behavioural and neurobiological measures, we conducted a PCA using individual loadings for the behavioural components (punished responding, CS+ aversion, unpunished responding, early context fear) and network components (N1, N2, N3) as input. Results were varimax-rotated, and the optimal solution was selected as described above. Relationships were then directly assessed using linear regression analyses in GraphPad Prism. Analyses were performed between each behavioural component (excl. CS+ aversion) and each network, and models were fit using the least squares regression method.

## References

Balleine, B. W., Delgado, M. R., & Hikosaka, O. (2007). The Role of the Dorsal Striatum in Reward and Decision-Making. Journal of Neuroscience, 27(31), 8161–8165. 10.1523/JNEUROSCI.1554-07.2007

Bankhead, P., Loughrey, M. B., Fernández, J. A., Dombrowski, Y., McArt, D. G., Dunne, P. D., McQuaid, S., Gray, R. T., Murray, L. J., Coleman, H. G., James, J. A., Salto-Tellez, M., & Hamilton, P. W. (2017). QuPath: Open source software for digital pathology image analysis. Scientific Reports, 7(1), 16878. 10.1038/s41598-017-17204-5

Bechara, A., Tranel, D., & Damasio, H. (2000). Characterization of the decision-making deficit of patients with ventromedial prefrontal cortex lesions. Brain, 123(11), 2189–2202. 10.1093/brain/123.11.2189

Bouton, M. E., & Broomer, M. C. (2023). Learning to stop responding. Behavioural Processes, 206, 104830. 10.1016/j.beproc.2023.104830

Broman-Fulks, J. J., Urbaniak, A., Bondy, C. L., & Toomey, K. J. (2014). Anxiety sensitivity and risk-taking behavior. Anxiety, Stress, & Coping, 27(6), 619–632. 10.1080/10615806.2014.896906

Broomer, M. C., & Bouton, M. E. (2024). Infralimbic cortex plays a similar role in the punishment and extinction of instrumental behavior. Neurobiology of Learning and Memory, 211, 107926. 10.1016/j.nlm.2024.107926

Burgos-Robles, A., Kimchi, E. Y., Izadmehr, E. M., Porzenheim, M. J., Ramos-Guasp, W. A., Nieh, E. H., Felix-Ortiz, A. C., Namburi, P., Leppla, C. A., Presbrey, K. N., Anandalingam, K. K., Pagan-Rivera, P. A., Anahtar, M., Beyeler, A., & Tye, K. M. (2017). Amygdala inputs to prefrontal cortex guide behavior amid conflicting cues of reward and punishment. Nature Neuroscience, 20(6), 824–835. 10.1038/nn.4553

Chen, B. T., Yau, H.-J., Hatch, C., Kusumoto-Yoshida, I., Cho, S. L., Hopf, F. W., & Bonci, A. (2013). Rescuing cocaine-induced prefrontal cortex hypoactivity prevents compulsive cocaine seeking. Nature, 496(7445), 359–362. 10.1038/nature12024

Chen, Y., Wang, G., Zhang, W., Han, Y., Zhang, L., Xu, H., Meng, S., Lu, L., Xue, Y., & Shi, J. (2022). An orbitofrontal cortex–anterior insular cortex circuit gates compulsive cocaine use. Science Advances, 8(51), eabq5745. 10.1126/sciadv.abq5745

Clark, L., Bechara, A., Damasio, H., Aitken, M. R. F., Sahakian, B. J., & Robbins, T. W. (2008). Differential effects of insular and ventromedial prefrontal cortex lesions on risky decision-making. Brain, 131(5), 1311–1322. 10.1093/brain/awn066

Corr, P. J. (2004). Reinforcement sensitivity theory and personality. Neuroscience & Biobehavioral Reviews, 28(3), 317–332. 10.1016/j.neubiorev.2004.01.005

Cox, J., & Witten, I. B. (2019). Striatal circuits for reward learning and decision-making. Nature Reviews Neuroscience, 20(8), 482–494. 10.1038/s41583-019-0189-2

Dadds, M. R., & Salmon, K. (2003). Punishment Insensitivity and Parenting: Temperament and Learning as Interacting Risks for Antisocial Behavior. Clinical Child and Family Psychology Review, 6(2), 69–86. 10.1023/A:1023762009877

Davis, M. (1992). The Role of the Amygdala in Fear and Anxiety. Annual Review of Neuroscience, 15(Volume 15, 1992), 353–375. 10.1146/annurev.ne.15.030192.002033

de Lima, M. A. X., Baldo, M. V. C., Oliveira, F. A., & Canteras, N. S. (2022). The anterior cingulate cortex and its role in controlling contextual fear memory to predatory threats. eLife, 11, e67007. 10.7554/eLife.67007

De Ruiter, M. B., Veltman, D. J., Goudriaan, A. E., Oosterlaan, J., Sjoerds, Z., & Van Den Brink, W. (2009). Response Perseveration and Ventral Prefrontal Sensitivity to Reward and Punishment in Male Problem Gamblers and Smokers. Neuropsychopharmacology, 34(4), 1027–1038. 10.1038/npp.2008.175

Domi, E., Xu, L., Toivainen, S., Nordeman, A., Gobbo, F., Venniro, M., Shaham, Y., Messing, R. O., Visser, E., Van Den Oever, M. C., Holm, L., Barbier, E., Augier, E., & Heilig, M. (2021). A neural substrate of compulsive alcohol use. Science Advances, 7(34), eabg9045. 10.1126/sciadv.abg9045

Elster, E. M., Pauli, R., Baumann, S., De Brito, S. A., Fairchild, G., Freitag, C. M., Konrad, K., Roessner, V., Brazil, I. A., Lockwood, P. L., & Kohls, G. (2024). Impaired Punishment Learning in Conduct Disorder. Journal of the American Academy of Child & Adolescent Psychiatry, 63(4), 454–463. 10.1016/j.jaac.2023.05.032

Engeln, M., & Ahmed, S. H. (2024). The multiple faces of footshock punishment in animal research on addiction. Neurobiology of Learning and Memory, 213, 107955. 10.1016/j.nlm.2024.107955

Eshel, N., & Roiser, J. P. (2010). Reward and Punishment Processing in Depression. Biological Psychiatry, 68(2), 118–124. 10.1016/j.biopsych.2010.01.027

Falconi-Sobrinho, L. L., Anjos-Garcia, T. D., Rebelo, M. A., Hernandes, P. M., Almada, R. C., Tanus-Santos, J. E., & Coimbra, N. C. (2024). The anterior cingulate cortex and its interface with the dorsal periaqueductal grey regulating nitric oxide-mediated panic-like behaviour and defensive antinociception. Neuropharmacology, 245, 109831. 10.1016/j.neuropharm.2023.109831

Falconi-Sobrinho, L. L., Dos Anjos-Garcia, T., & Coimbra, N. C. (2021). Nitric oxide-mediated defensive and antinociceptive responses organised at the anterior hypothalamus of mice are modulated by glutamatergic inputs from area 24b of the cingulate cortex. Journal of Psychopharmacology, 35(1), 78–90. 10.1177/0269881120967881

Falconi-Sobrinho, L. L., Dos Anjos-Garcia, T., Elias-Filho, D. H., & Coimbra, N. C. (2017). Unravelling cortico-hypothalamic pathways regulating unconditioned fear-induced antinociception and defensive behaviours. Neuropharmacology, 113, 367–385. 10.1016/j.neuropharm.2016.10.001

Fellows, L. K., & Farah, M. J. (2003). Ventromedial frontal cortex mediates affective shifting in humans: Evidence from a reversal learning paradigm. Brain, 126(8), 1830–1837. 10.1093/brain/awg180

Figee, M., Pattij, T., Willuhn, I., Luigjes, J., Van Den Brink, W., Goudriaan, A., Potenza, M. N., Robbins, T. W., & Denys, D. (2016). Compulsivity in obsessive–compulsive disorder and addictions. European Neuropsychopharmacology, 26(5), 856–868. 10.1016/j.euroneuro.2015.12.003

Freeman, L. C. (1977). A Set of Measures of Centrality Based on Betweenness. Sociometry, 40(1), 35–41. 10.2307/3033543

Freeman, L. C. (1978). Centrality in social networks conceptual clarification. Social Networks, 1(3), 215–239. 10.1016/0378-8733(78)90021-7

Gaetani, K., Jean-Richard-dit-Bressel, P., & McNally, G. P. (2025). Early contingency information enhances human punishment sensitivity when punishment is frequent but not rare. Behavioral Neuroscience, 139(4–5), 216–228. 10.1037/bne0000627

Giorgetta, C., Grecucci, A., Zuanon, S., Perini, L., Balestrieri, M., Bonini, N., Sanfey, A., & Brambilla, P. (2012). Reduced Risk-Taking Behavior as a Trait Feature of Anxiety. Emotion, 12(6), 1373–1383. 10.1037/a0029119

Giustino, T. F., & Maren, S. (2015). The Role of the Medial Prefrontal Cortex in the Conditioning and Extinction of Fear. Frontiers in Behavioral Neuroscience, 9. 10.3389/fnbeh.2015.00298

Goldsborough, T., O’Callaghan, A., Inglis, F., Leplat, L., Filby, A., Bilen, H., & Bankhead, P. (2024). A novel channel invariant architecture for the segmentation of cells and nuclei in multiplexed images using InstanSeg (p. 2024.09.04.611150). bioRxiv. 10.1101/2024.09.04.611150

Grewe, B. F., Gründemann, J., Kitch, L. J., Lecoq, J. A., Parker, J. G., Marshall, J. D., Larkin, M. C., Jercog, P. E., Grenier, F., Li, J. Z., Lüthi, A., & Schnitzer, M. J. (2017). Neural ensemble dynamics underlying a long-term associative memory. Nature, 543(7647), 670–675. 10.1038/nature21682

Groeneboom, N. E., Yates, S. C., Puchades, M. A., & Bjaalie, J. G. (2020). Nutil: A Pre- and Post-processing Toolbox for Histological Rodent Brain Section Images. Frontiers in Neuroinformatics, 14. 10.3389/fninf.2020.00037

Guimerà, R., & Nunes Amaral, L. A. (2005). Functional cartography of complex metabolic networks. Nature, 433(7028), 895–900. 10.1038/nature03288

Hevey, D., Thomas, K., Laureano-Schelten, S., Looney, K., & Booth, R. (2017). Clinical Depression and Punishment Sensitivity on the BART. Frontiers in Psychology, 8. 10.3389/fpsyg.2017.00670

Ishikawa, J., Sakurai, Y., Ishikawa, A., & Mitsushima, D. (2020). Contribution of the prefrontal cortex and basolateral amygdala to behavioral decision-making under reward/punishment conflict. Psychopharmacology, 237(3), 639–654. 10.1007/s00213-019-05398-7

Jean-Richard-dit-Bressel, P., Gaetani, K., Zeng, L., Weidemann, G., & McNally, G. P. (2024). Translational research in punishment learning. Behavioral Neuroscience, 138(3), 143–151. 10.1037/bne0000587

Jean-Richard-Dit-Bressel, P., Killcross, S., & McNally, G. P. (2018). Behavioral and neurobiological mechanisms of punishment: Implications for psychiatric disorders. Neuropsychopharmacology, 43(8), 1639–1650. 10.1038/s41386-018-0047-3

Jean-Richard-dit-Bressel, P., Lee, J. C., Liew, S. X., Weidemann, G., Lovibond, P. F., & McNally, G. P. (2021). Punishment insensitivity in humans is due to failures in instrumental contingency learning. eLife, 10, e69594. 10.7554/eLife.69594

Jean-Richard-dit-Bressel, P., Lee, J. C., Liew, S. X., Weidemann, G., Lovibond, P. F., & McNally, G. P. (2023). A cognitive pathway to punishment insensitivity. Proceedings of the National Academy of Sciences, 120(15), e2221634120. 10.1073/pnas.2221634120

Jean-Richard-dit-Bressel, P., Ma, C., Bradfield, L. A., Killcross, S., & McNally, G. P. (2019). Punishment insensitivity emerges from impaired contingency detection, not aversion insensitivity or reward dominance. eLife, 8, e52765. 10.7554/eLife.52765

Jean-Richard-dit-Bressel, P., & McNally, G. P. (2016). Lateral, not medial, prefrontal cortex contributes to punishment and aversive instrumental learning. Learning & Memory, 23(11), 607–617. 10.1101/lm.042820.116

Killcross, A. S., Everitt, B. J., & Robbins, T. W. (1997). Symmetrical effects of amphetamine and alpha-flupenthixol on conditioned punishment and conditioned reinforcement: Contrasts with midazolam. Psychopharmacology, 129(2), 141–152. 10.1007/s002130050174

Kimbrough, A., Lurie, D. J., Collazo, A., Kreifeldt, M., Sidhu, H., Macedo, G. C., D’Esposito, M., Contet, C., & George, O. (2020). Brain-wide functional architecture remodeling by alcohol dependence and abstinence. Proceedings of the National Academy of Sciences, 117(4), 2149–2159. 10.1073/pnas.1909915117

Kintali, S. (2008). Betweenness Centrality: Algorithms and Lower Bounds (No. 0809.1906). arXiv. 10.48550/arXiv.0809.1906

Kleven, H., Bjerke, I. E., Clascá, F., Groenewegen, H. J., Bjaalie, J. G., & Leergaard, T. B. (2023). Waxholm Space atlas of the rat brain: A 3D atlas supporting data analysis and integration. Nature Methods, 20(11), 1822–1829. 10.1038/s41592-023-02034-3

Liley, A. E., Gabriel, D. B. K., Sable, H. J., & Simon, N. W. (2019). Sex Differences and Effects of Predictive Cues on Delayed Punishment Discounting. Eneuro, 6(4), ENEURO.0225-19.2019. 10.1523/ENEURO.0225-19.2019

Limpens, J. H. W., Damsteegt, R., Broekhoven, M. H., Voorn, P., & Vanderschuren, L. J. M. J. (2015). Pharmacological inactivation of the prelimbic cortex emulates compulsive reward seeking in rats. Brain Research, 1628, 210–218. 10.1016/j.brainres.2014.10.045

Ma, C., Jean-Richard-dit-Bressel, P., Roughley, S., Vissel, B., Balleine, B. W., Killcross, S., & Bradfield, L. A. (2020). Medial Orbitofrontal Cortex Regulates Instrumental Conditioned Punishment, but not Pavlovian Conditioned Fear. Cerebral Cortex Communications, 1(1), tgaa039. 10.1093/texcom/tgaa039

Maier, S. F., & Jackson, R. L. (1979). Learned Helplessness: All of us were Right (And Wrong): Inescapable Shock has Multiple Effects1. In G. H. Bower (Ed.), Psychology of Learning and Motivation (Vol. 13, pp. p155–218). Academic Press. 10.1016/S0079-7421(08)60083-3

Maier, S. F., & Seligman, M. E. P. (2016). Learned Helplessness at Fifty: Insights from Neuroscience. Psychological Review, 123(4), 349–367. 10.1037/rev0000033

Marchant, N. J., Campbell, E. J., & Kaganovsky, K. (2018). Punishment of alcohol-reinforced responding in alcohol preferring P rats reveals a bimodal population: Implications for models of compulsive drug seeking. Progress in Neuro-Psychopharmacology and Biological Psychiatry, 87, 68–77. 10.1016/j.pnpbp.2017.07.020

Maren, S., & Quirk, G. J. (2004). Neuronal signalling of fear memory. Nature Reviews Neuroscience, 5(11), 844–852. 10.1038/nrn1535

McDonald, A. J., Nemat, P., van ‘t Hullenaar, T., Schetters, D., van Mourik, Y., Alonso-Lozares, I., De Vries, T. J., & Marchant, N. J. (2024). Punishment-resistant alcohol intake is mediated by the nucleus accumbens shell in female rats. Neuropsychopharmacology, 49(13), 2022– 2031. 10.1038/s41386-024-01940-0

McNally, G. P., Jean-Richard-dit-Bressel, P., Millan, E. Z., & Lawrence, A. J. (2023). Pathways to the persistence of drug use despite its adverse consequences. Molecular Psychiatry, 28(6), 2228–2237. 10.1038/s41380-023-02040-z

O’Brien, B. S., & Frick, P. J. (1996). Reward dominance: Associations with anxiety, conduct problems, and psychopathy in children. Journal of Abnormal Child Psychology, 24(2), 223– 240. 10.1007/BF01441486

Orsini, C. A., Trotta, R. T., Bizon, J. L., & Setlow, B. (2015). Dissociable Roles for the Basolateral Amygdala and Orbitofrontal Cortex in Decision-Making under Risk of Punishment. The Journal of Neuroscience, 35(4), 1368–1379. 10.1523/JNEUROSCI.3586-14.2015

Palminteri, S., Clair, A.-H., Mallet, L., & Pessiglione, M. (2012). Similar Improvement of Reward and Punishment Learning by Serotonin Reuptake Inhibitors in Obsessive-Compulsive Disorder. Biological Psychiatry, 72(3), 244–250. 10.1016/j.biopsych.2011.12.028

Pelloux, Y., Everitt, B. J., & Dickinson, A. (2007). Compulsive drug seeking by rats under punishment: Effects of drug taking history. Psychopharmacology, 194(1), 127–137. 10.1007/s00213-007-0805-0

Pelloux, Y., Murray, J. E., & Everitt, B. J. (2013). Differential roles of the prefrontal cortical subregions and basolateral amygdala in compulsive cocaine seeking and relapse after voluntary abstinence in rats. European Journal of Neuroscience, 38(7), 3018–3026. 10.1111/ejn.12289

Piantadosi, P. T., Yeates, D. C. M., Wilkins, M., & Floresco, S. B. (2017). Contributions of basolateral amygdala and nucleus accumbens subregions to mediating motivational conflict during punished reward-seeking. Neurobiology of Learning and Memory, 140, 92–105. 10.1016/j.nlm.2017.02.017

Puchades, M. A., Csucs, G., Ledergerber, D., Leergaard, T. B., & Bjaalie, J. G. (2019). Spatial registration of serial microscopic brain images to three-dimensional reference atlases with the QuickNII tool. PLOS ONE, 14(5), e0216796. 10.1371/journal.pone.0216796

Reineberg, A. E., Andrews-Hanna, J. R., Depue, B. E., Friedman, N. P., & Banich, M. T. (2015). Resting-state networks predict individual differences in common and specific aspects of executive function. NeuroImage, 104, 69–78. 10.1016/j.neuroimage.2014.09.045

Resstel, L. B. M., Souza, R. F., & Guimarães, F. S. (2008). Anxiolytic-like effects induced by medial prefrontal cortex inhibition in rats submitted to the Vogel conflict test. Physiology & Behavior, 93(1–2), 200–205. 10.1016/j.physbeh.2007.08.009

Rich, A. S., & Gureckis, T. M. (2018). The limits of learning: Exploration, generalization, and the development of learning traps. Journal of Experimental Psychology: General, 147(11), 1553–1570. 10.1037/xge0000466

Robinson, T. E., & Berridge, K. C. (2003). Addiction. Annual Review of Psychology, 54(1), 25–53. 10.1146/annurev.psych.54.101601.145237

Rubinov, M., & Sporns, O. (2010). Complex network measures of brain connectivity: Uses and interpretations. NeuroImage, 52(3), 1059–1069. 10.1016/j.neuroimage.2009.10.003

Sah, P., Faber, E. S. L., Lopez De Armentia, M., & Power, J. (2003). The Amygdaloid Complex: Anatomy and Physiology. Physiological Reviews, 83(3), 803–834. 10.1152/physrev.00002.2003

Schindelin, J., Arganda-Carreras, I., Frise, E., Kaynig, V., Longair, M., Pietzsch, T., Preibisch, S., Rueden, C., Saalfeld, S., Schmid, B., Tinevez, J.-Y., White, D. J., Hartenstein, V., Eliceiri, K., Tomancak, P., & Cardona, A. (2012). Fiji: An open-source platform for biological-image analysis. Nature Methods, 9(7), 676–682. 10.1038/nmeth.2019

Seligman, M. E. (1972). Learned helplessness. Annual Review of Medicine, 23, 407–412. 10.1146/annurev.me.23.020172.002203

Sengupta, A., Yau, J. O. Y., Jean-Richard-Dit-Bressel, P., Liu, Y., Millan, E. Z., Power, J. M., & McNally, G. P. (2018). Basolateral Amygdala Neurons Maintain Aversive Emotional Salience. The Journal of Neuroscience, 38(12), 3001–3012. 10.1523/JNEUROSCI.2460-17.2017

Smith, S. M., Fox, P. T., Miller, K. L., Glahn, D. C., Fox, P. M., Mackay, C. E., Filippini, N., Watkins, K. E., Toro, R., Laird, A. R., & Beckmann, C. F. (2009). Correspondence of the brain’s functional architecture during activation and rest. Proceedings of the National Academy of Sciences, 106(31), 13040–13045. 10.1073/pnas.0905267106

Sporns, O., Honey, C. J., & Kötter, R. (2007). Identification and Classification of Hubs in Brain Networks. PLOS ONE, 2(10), e1049. 10.1371/journal.pone.0001049

Terstege, D. J., & Epp, J. R. (2023). Network Neuroscience Untethered: Brain-Wide Immediate Early Gene Expression for the Analysis of Functional Connectivity in Freely Behaving Animals. Biology, 12(1), 34. 10.3390/biology12010034

Thorndike, E. L. (1911). Animal intelligence: Experimental studies (pp. viii, 297). Macmillan Press. 10.5962/bhl.title.55072

Torres, O. V., Jayanthi, S., Ladenheim, B., McCoy, M. T., Krasnova, I. N., & Cadet, J. L. (2017). Compulsive methamphetamine taking under punishment is associated with greater cue-induced drug seeking in rats. Behavioural Brain Research, 326, 265–271. 10.1016/j.bbr.2017.03.009

Vanderschuren, L. J., Minnaard, A. M., Smeets, J. A., & Lesscher, H. M. (2017). Punishment models of addictive behavior. Current Opinion in Behavioral Sciences, 13, 77–84. 10.1016/j.cobeha.2016.10.007

Verharen, J. P. H., Heuvel, M. W. van den, Luijendijk, M., Vanderschuren, L. J. M. J., & Adan, R. A. H. (2019). Corticolimbic Mechanisms of Behavioral Inhibition under Threat of Punishment. Journal of Neuroscience, 39(22), 4353–4364. 10.1523/JNEUROSCI.2814-18.2019

Vetere, G., Kenney, J. W., Tran, L. M., Xia, F., Steadman, P. E., Parkinson, J., Josselyn, S. A., & Frankland, P. W. (2017). Chemogenetic Interrogation of a Brain-wide Fear Memory Network in Mice. Neuron, 94(2), 363-374.e4. 10.1016/j.neuron.2017.03.037

Wheeler, A. L., Teixeira, C. M., Wang, A. H., Xiong, X., Kovacevic, N., Lerch, J. P., McIntosh, A. R., Parkinson, J., & Frankland, P. W. (2013). Identification of a Functional Connectome for Long-Term Fear Memory in Mice. PLOS Computational Biology, 9(1), e1002853. 10.1371/journal.pcbi.1002853

Yates, S. C., Groeneboom, N. E., Coello, C., Lichtenthaler, S. F., Kuhn, P.-H., Demuth, H.-U., Hartlage-Rübsamen, M., Roßner, S., Leergaard, T., Kreshuk, A., Puchades, M. A., & Bjaalie, J. G. (2019). QUINT: Workflow for Quantification and Spatial Analysis of Features in Histological Images From Rodent Brain. Frontiers in Neuroinformatics, 13. 10.3389/fninf.2019.00075

Zeng, L., Park, H. R. P., McNally, G. P., & Jean-Richard-dit-Bressel, P. (2025). Causal inference and cognitive-behavioral integration deficits drive stable variation in human punishment sensitivity. Communications Psychology, 3(1), 103. 10.1038/s44271-025-00284-9

