## Supplemental Materials for "Relationships between phenotypic differences in punishment behaviour and prefrontal cortex network engagement"

**Supplementary Materials**

**- Figure S1**

**- DREADD manipulation: Methods, Results, Figures S2-4**

**Cluster Solutions**

A

| Clusters ($k$) | *n* of smallest cluster | Min | Max | Mean | SD |
|---|---|---|---|---|---|
| **2** | 32 | .040 | .517 | .344 | .131 |
| **3** | 10 | -.007# | .511 | .297 | .126 |
| **4** | 10 | .036 | .507 | .227 | .119 |
| **5** | 1# | -.016# | 1.000 | .248 | .147 |
| **6** | 1# | -.005# | 1.000 | .251 | .151 |
| **7** | 1# | .038 | 1.000 | .255 | .143 |

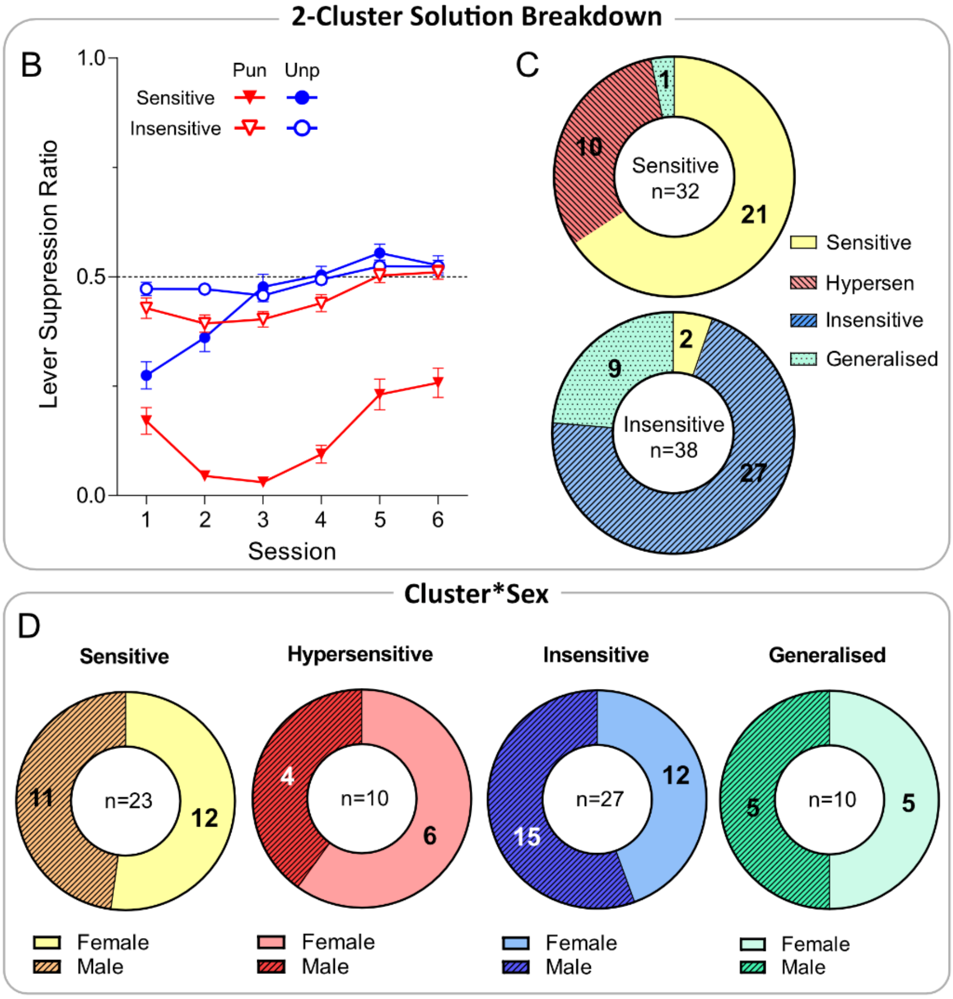


**Figure S1. [A]** Descriptives for k-means cluster solutions. Table shows number of subjects in smallest cluster, minimum silhouette value, maximum silhouette value, mean silhouette value, and standard deviation of silhouette values per cluster solution. Cluster solutions with single-subject clusters and/or negative silhouette values ($k$ = 3,5,6,7) were deemed suboptimal. **[B]** Mean ± SEM lever suppression ratios per lever (punished R1 [Pun], unpunished R2 [Unp]) by $k$=2 clusters (sensitive, insensitive). **[C]** Relation between $k$=2 clusters (sensitive, insensitive) and $k$=4 clusters assignments. Each $k$=4 cluster almost exclusively derived from one of the two $k$=2 clusters, indicating subclusters of sensitive vs. insensitive phenotypes. **[D]** Breakdown of sex per $k$=4 cluster.

**DREADD manipulation: Methods & Results**

**Methods**

***Surgery***

All rats underwent stereotaxic surgery to express excitatory hm3D DREADD in mOFC. Rats were anesthetized with isoflurane (5% induction; 2% maintenance) mixed with oxygen and placed in a stereotaxic frame (David Kopf Instruments). Prior to incision, rats received the analgesic carprofen (Rimadyl, Zoetis; 5 mg/kg) via a subcutaneous injection and 0.5% bupivacaine under the incision site. An incision was performed to expose the skull and a bilateral craniotomy was performed above mOFC. 750nL of AAV-CAMKII-hM3Dq-MCITRINE (Addgene, MA, USA) was injected bilaterally into the mOFC (AP: +4.2, ML: ±0.5, DV: -4.8mm from bregma) via 23-gauge 5 mL Hamilton syringe at a rate of 250nL/min. The syringe was left in place for 5 min to allow for virus diffusion. The incision was sutured, and antibiotic (Duplocillin, Intervet; 40,000-60,000 IU/kg) was given intraperitoneally.

Rats were given 3 weeks for post-operative recovery and expression of hm3D.

***hM3D ligand***

hM3D activation was attempted using systemic (i.p.) injections of hM3D ligands deschloroclozapine (DCZ; 0.1mg/mL/kg) or clozapine (CLZ; 0.1mg/mL/kg). Each rat was either exposed to DCZ ($N$ = 23) or CLZ ($N$ = 47) across the experiment. No differences between DCZ or CLZ were observed; DCZ and CLZ are reported in aggregate as "Ligand". Saline (1mL/kg) was used as a control comparison for Ligand.

***Injection procedure***

Rats received injections of saline and/or ligand at 3 points in the experiment. To habituate animals to the injection procedure, rats received dummy injections with an empty needle on the last 2 days of lever-press training.

To examine the effects of mOFC activation on acquisition of aversion, rats received injections 15mins before the first 2 sessions of conditioned punishment (punishment acquisition). Rats were randomly assigned to receive 2 sessions of saline or 2 sessions of ligand (between-subjects).

To examine the effects of mOFC activation on expression of learned behaviour, rats received injections 15mins before sessions 7 and 9 of conditioned punishment (punishment expression). Each rat received saline or ligand injection prior to expression sessions (within-subjects, order counterbalanced). Sessions before and after each injection session was injection-free to limit carry-over effects and provide a measure of behaviour before and after injections.

To verify the efficacy of the DREADD manipulation, rats received either saline or ligand injections 2hrs prior to perfusions to allow for DREADD activation and cFos induction.

***Immunohistochemistry***

For the first batch ($n$=23), brain tissue was incubated in Triton X-100 solution for 2hrs (0.5% Triton X-100, 5% normal donkey serum [NDS], 5% normal goat serum [NGS] in 0.1M PBS) and incubated overnight in primary antibody solution (1:1000 chicken anti-GFP, 1:1000 rabbit anti-cFos, 0.2% Triton X-100, 1% NDS, 1% NGS in 0.1M phosphate buffer solution [PBS]). Unbound antibodies were washed off in PBS (3 times, 10mins each) and incubated in secondary antibody solution (1:500 AF488 anti-chicken, 1:500 AF594 donkey anti-rabbit, 0.2% Triton X-100, 1% NDS, 1% NGS in PBS) for 6hrs.

For the second ($n$=23) and third batches ($n$=23), brain tissue was incubated in Trixon X-100 for 1hr (0.5% Triton X-100, 10% NDS, 10% NGS in 0.1M PBS) and incubated overnight in primary antibody solution (1:1000 rabbit anti-GFP, 1:3000 guinea pig anti-cFos, 0.5% Triton X-100, 2% NDS, 2% NGS in PBS). Unbound antibodies were washed off in PBS (3 times, 10mins each) and incubated in secondary antibody solution (1:500 AF488 donkey anti-rabbit, 1:500 AF647 goat anti-guinea pig, 0.2% Triton X-100, 1% NDS, 1% NGS in PBS) for 2hrs.

### Results

Animals with <10 hM3D+ cells/mm$^2$ in mOFC were considered as not having sufficient mOFC hM3D expression, and were thus excluded from all DREADD-related analyses ($n$=5 [saline], $n$=1 [DCZ], $n$=5 [CLZ]), leaving $N$=59.

#### *Punishment acquisition*

Ligand injections on the first 2 days of conditioned punishment had no effect on acquisition of punishment avoidance . There were no significant differences between ligand and saline groups in lever suppression (Group: $F_{(1,57)}$=.429, $p$=.515; Group*Lever: $F_{(1,57)}$=.042, $p$=.838) (**Figure S2A**) or CS suppression (Group: $F_{(1,57)}$=.214, $p$=.646; Group*CS: $F_{(1,57)}$=3.089, $p$=.084) (**Figure S2B**). Saline and ligand groups did not significantly differ in punishment-sensitivity cluster assignment ($\chi^2_{(3)}$=2.698, $p$=.441) (**Figure S2C**).

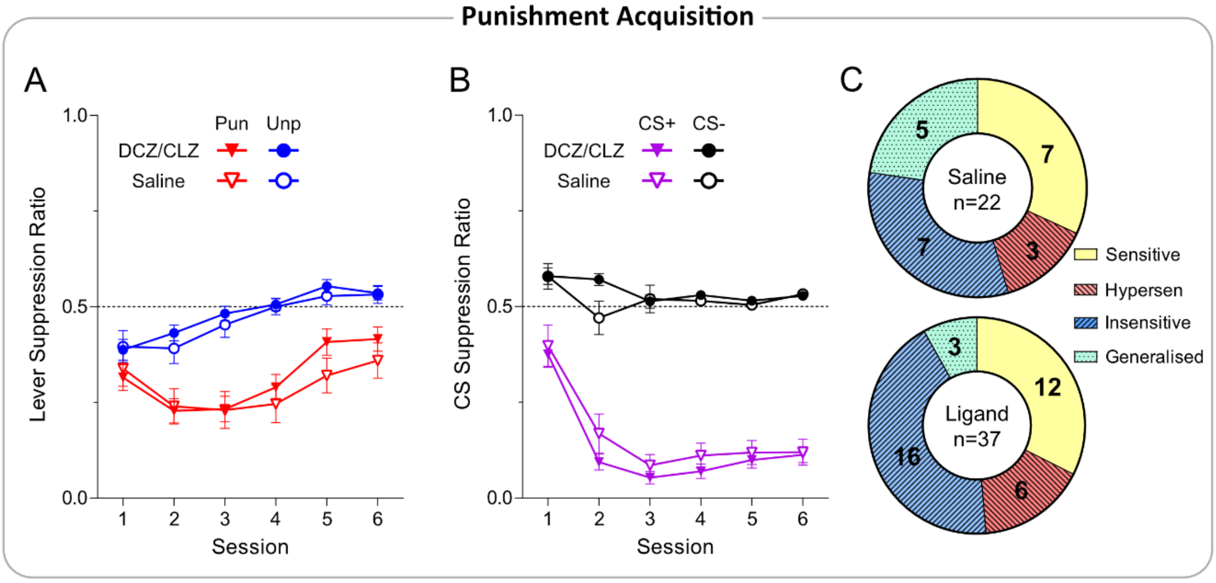


**Figure S2. Ligand injections had no effect on punishment acquisition. [A]** Mean ± SEM lever suppression across punishment acquisition sessions by acquisition injection condition (Saline vs. Ligand [between-subjects]). No effect of DREADD ligand was observed. **[B]** Mean ± SEM CS suppression across punishment acquisition sessions by acquisition injection condition (Saline vs. Ligand [between-subjects]). No effect of DREADD ligand was observed. **[C]** Breakdown of cluster assignment by injection condition (Saline vs. Ligand [between-subjects]).

#### *Punishment expression*

Ligand efficacy on punishment expression was assessed within-subjects, where rats received saline or ligand across 2 sessions in late-punishment. Ligand injections had no effect on expression of punishment avoidance. During injection sessions, rats continued to suppress punished over unpunished responding (Lever: $F_{(1,58)}$=20.65, $p$<.001, $\eta^2$=.263) irrespective of injection type (Drug: $F_{(1,58)}$=.010, $p$=.919; Drug *Lever: $F_{(1,57)}$=.981, $p$=.326) (**Figure S3A**). Rats continued to suppress more during CS+ than CS- presentations (CS: $F_{(1,58)}$=282.6, $p$<.001, $\eta^2$=.830), and expression injection had no effect on this (Drug: $F_{(1,58)}$=3.107, $p$=.083; Drug *CS: $F_{(1,58)}$=.101, $p$=.751) (**Figure S3B**).

There were also no significant interaction between punishment-sensitivity clusters and drug for lever suppression (Ligand*Cluster: $F_{(3,55)}$=.406, $p$=.749) and CS suppression (Ligand*Cluster: $F_{(3,55)}$=.375, $p$=.771) during expression sessions.

Together, these results indicate the ligand had no effects on punishment expression.

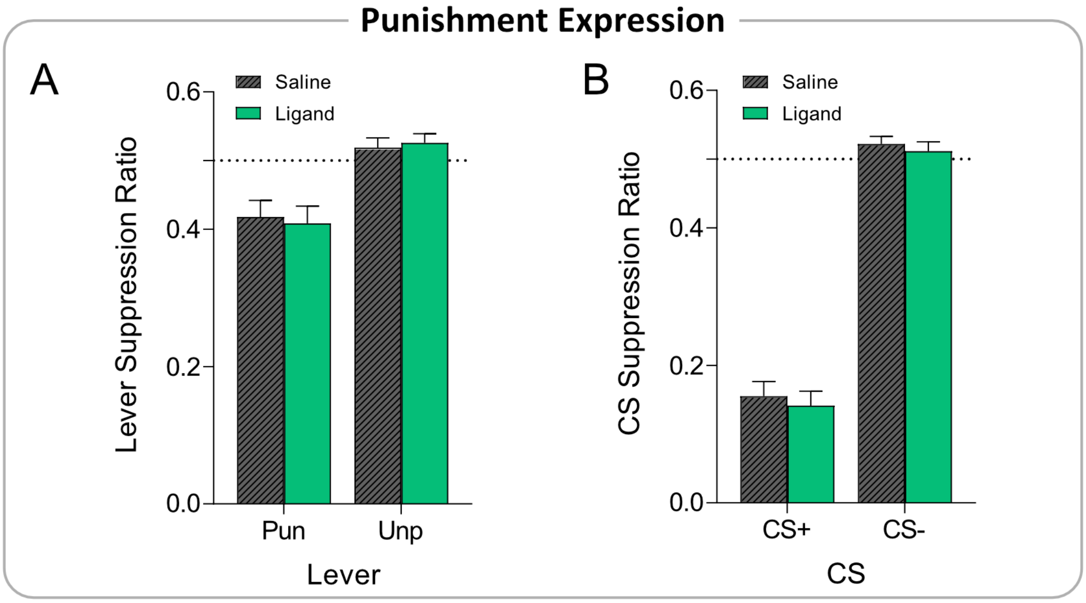


**Figure S3. Effect of ligand injection on punishment expression. [A]** Mean ± SEM lever suppression during punishment expression session, by injection condition (Saline vs. Ligand [within-subjects]). No effect of DREADD ligand was observed. **[B]** Mean ± SEM CS suppression during punishment expression session, by injection condition (Saline vs. Ligand [within-subjects]). No effect of DREADD ligand was observed.

***c-Fos expression***

To verify the efficacy of DREADD manipulation, we assessed the effect of perfusion ligand on c-Fos expression. If ligands successfully activated hM3Dq-expressing cells, rats receiving ligands prior to perfusion should show higher c-Fos densities (particularly in DREADD-targeted mOFC), and greater c-Fos in hM3Dq-expressing cells, relative to saline-injected animals. There was no significant effect of injections on overall c-Fos in mOFC ($F_{(1,57)}$=.535, $p$=.468) or any other PFC region (Max $F$-value [FrA]: $F_{(1,52)}$=1.806, $p$=.185) (**Figure S4A**). Similarly, perfusion injections did not significantly affect c-Fos levels in hM3Dq-expressing cells ($F_{(1,57)}$=.080, $p$=.779) (**Figure S4B**). This strongly suggests the DREADD manipulation failed to activate mOFC as intended.

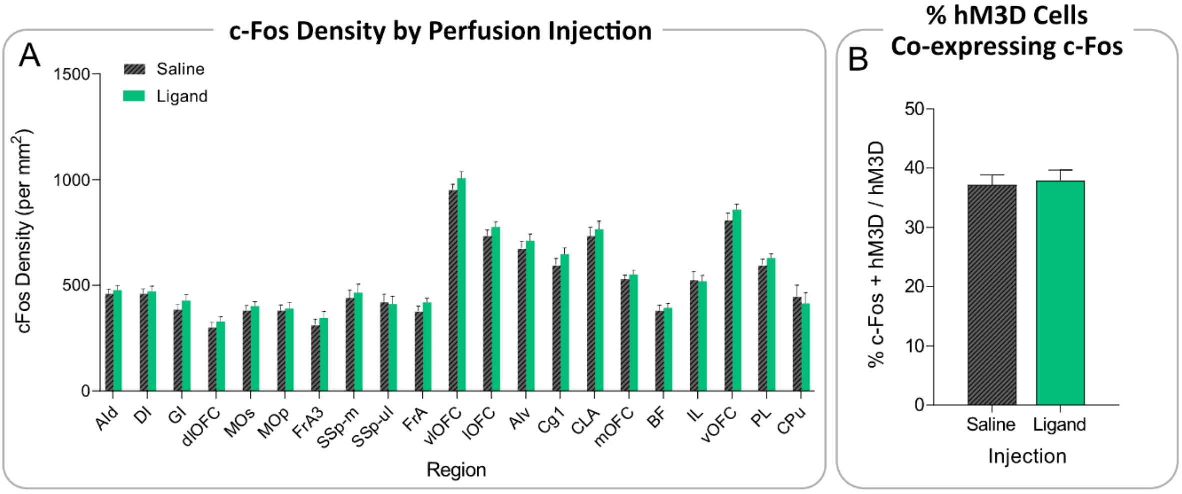


**Figure S4. Ligand injections failed to activate hM3D cells. [A]** Mean ± SEM c-Fos expression across cortical regions by perfusion injection condition (Saline vs. Ligand). No effect of DREADD ligand was observed. **[B]** Mean ± SEM c-Fos expression in hm3D-expressing cells by perfusion injection condition (Saline vs. Ligand). No effect of DREADD ligand was observed.
